# Uncovering the regulatory network of the small RNA SuhB and its contribution to stress resistance in *Sphingopyxis granuli* TFA

**DOI:** 10.64898/2025.12.09.693200

**Authors:** Inmaculada García-Romero, Alberto Pires-Acosta, Belén Floriano, Francisca Reyes-Ramírez

## Abstract

Post-transcriptional regulation by small RNAs (sRNAs) enables bacteria to fine-tune gene expression and rapidly adapt to fluctuating environmental conditions. In *Sphingopyxis granuli* TFA, SuhB, the only sRNA characterized to date in this strain, was previously shown to repress *thnR* translation to control tetralin degradation under carbon catabolite repression conditions. Here, we reveal additional regulatory roles of SuhB beyond carbon metabolism. Deletion of *suhB* increases sensitivity to diverse abiotic stresses, including osmotic, oxidative, desiccation, and copper stress. Label-free quantitative proteomic analysis indicates widespread alterations in the proteome in the absence of SuhB, affecting metabolic pathways and membrane-associated processes. Moreover, a LysR-type transcription factor mutant, identified as a direct activator of *suhB,* shows similar phenotypes. Together, these findings demonstrate that SuhB functions as a global post-transcriptional regulator, coordinating metabolic balance, membrane composition, and stress resistance in *S. granuli* TFA, highlighting the critical role of sRNA-mediated regulation in environmental bacteria.

## Introduction

Gene expression in bacteria is orchestrated by a diverse array of mechanisms operating at multiple levels. While transcriptional control initiation dictates the initial production of RNA transcripts, post-transcriptional regulation adds an additional layer of modulation controlling the translation of the messenger RNA. These layers of regulation allow bacteria to rapidly adapt to fluctuating environmental conditions and optimize resource utilization, which is crucial for survival in dynamic or stressful habitats. In prokaryotes, post-transcriptional regulation is predominantly mediated by two key players: non-coding RNAs (ncRNAs) and RNA-binding proteins (RBPs) (1, 2). Among bacterial ncRNAs, and in contrast to antisense RNAs and other cis-encoded regulatory RNAs, trans-encoded small RNAs (sRNAs) are found in genomic regions separate from their target genes. While antisense sRNAs generally exhibit extensive complementarity and regulate a single overlapping mRNA, trans-encoded sRNAs share only limited complementarity with their targets and often require RNA chaperones such as Hfq to facilitate base-pairing, enabling them to control multiple genes and have broader regulatory networks (*3*). The primary mode of action of trans-encoded sRNAs is through imperfect base-pairing with target mRNAs, which can interfere with translation and/or affect RNA stability, most commonly by binding near the ribosome binding site (RBS) (*4*). However, trans-encoded sRNAs can also act at more distal regions of the 5′-untranslated region (5′-UTR) upstream of the RBS, or even within the open reading frame (*5*). Through these mechanisms, sRNAs can orchestrate complex regulatory networks that influence diverse cellular processes. Since the discovery of the sRNAs, multiple biological functions in bacteria have been described to be somehow regulated by them, such as virulence and host-pathogen interaction in the case of pathogenic bacteria (*6*), coordination of the stress responses by modulating the translation of key regulators such as *rpoS* in *Escherichia coli* (*7–9*), metabolism, such as SgrS in *E. coli*, which mediates the glucose-phosphate stress response (*10*) or antibiotic resistance (*11–13*).

While many studies have addressed the role of sRNAs in model and pathogenic bacteria, i.e. *Escherichia coli* and *Salmonella*, much less is known about their function in environmental species (*4*). Understanding sRNA-mediated regulation in these organisms is particularly relevant because it may reveal novel adaptative strategies for coping with fluctuating environmental conditions. *Sphingopyxis granuli* TFA is an alpha-proteobacterium isolated from the mud of the Rhine River (*14*) and able to degrade tetralin, an organic solvent (*15, 16*).

TFA is the first facultative anaerobe described within its genus, a feature that may allow this strain to invade anoxic and microoxic niches (*17*). Like other soil and aquatic bacteria, TFA continuously adjusts its physiology and metabolism to the variable environmental conditions, a process mediated by diverse mechanisms, including the activation of the General Stress Response (GSR). The GSR is a conserved regulatory response across many bacterial species, governing adaptation to diverse stresses such as desiccation, oxidative damage, nutrient limitation, and osmotic or envelope stress (*18–21*). In alpha-proteobacteria, the core of the response relies on a three-component regulatory system formed by the ECF sigma factor EcfG, the anti-sigma factor NepR, and the response regulator PhyR. Together, these elements control the transcriptional activation of stress related genes in response to environmental stress signals that are detected and transduced by histidine kinases, which regulate the phosphorylation status of PhyR (*18*). In TFA, the main components of the GSR regulatory cascade have been characterized, including two ECF sigma factors (EcfG1 and EcfG2), two anti-sigma factors (NepR1 and NepR2), and two anti-anti-sigma factors or response regulators (PhyR1 and PhyR2), thereby elucidating the specific role of each paralog in the activation and control of GSR transcriptional output under different stress conditions (*21–23*). The GSR regulon has also been characterized in TFA, and includes genes involved in stress adaptation, such as catalases, toxin–antitoxin systems, RND efflux components, and envelope-associated proteins (*23*).

Identification and characterization of sRNAs in alpha-proteobacteria has been carried out mainly in the photosynthetic *Rhodobacter sphaeroides* and in plant symbionts such as *Sinorhizobium meliloti* and *Bradyrhizobium japonicum* (*24–26*), although little is known about their broader regulatory networks. More recently, significant progress has also been made in elucidating sRNA-mediated regulation in *Caulobacter crescentus* (*27–29*).

In TFA, SuhB is the only small RNA characterized to date. It is a *trans*-encoded sRNA homologous to MmgR (make more granules regulator) described in *Sinorhizobium meliloti*. In this bacterium, MmgR plays a role in controlling polyhydroxybutyrate (PHB) accumulation. Deletion of *mmgR* leads to increased cellular PHB levels, suggesting that this sRNA may in some way influence the carbon flux toward PHB accumulation (*30*). In TFA, SuhB regulates the expression of tetralin degradation genes (*thn* genes) by inhibiting the translation of *thnR*, a LysR-type transcriptional activator of *thn* genes, under carbon catabolite repression (CCR) conditions or in the absence of tetralin. This repression is mediated through SuhB binding to the Shine Dalgarno region of the *thnR* mRNA, an interaction facilitated by the RNA chaperone Hfq (*16, 31*). Thus, a SuhB deletion mutant exhibits partial de-repression of *thn* genes in the presence of preferential carbon sources, because of the translation inhibition release of the *thnR* mRNA.

In this study, we investigate additional regulatory roles of SuhB in TFA and our results indicate that SuhB functions as a broader regulatory element. Comparative proteomic analyses performed under stress (stationary phase) and non-stress (exponential phase) conditions revealed a set of proteins whose abundance depends on SuhB, either directly or indirectly. Furthermore, cells lacking SuhB show reduced tolerance to several environmental stresses, suggesting that the changes in the proteome are necessary for survival of TFA under stress conditions. These findings suggest that SuhB functions as a global post-transcriptional regulator influencing multiple cellular processes.

We further identified a LysR-type transcriptional regulator that activates SuhB expression, and whose absence leads to reduced SuhB levels and decreased stress tolerance. This relationship supports a connection between the transcriptional control of SuhB and its regulatory function in TFA. Finally, some of the newly identified SuhB targets were experimentally confirmed, providing a more comprehensive view of the regulatory function of this sRNA in TFA.

## Materials and methods

### Media and growth conditions

*Escherichia coli* strains were cultured in Luria-Bertani (LB) medium or in M63 minimal medium (*32*) supplemented with 0.2% glycerol as the carbon source, at either 37 °C or 30 °C depending on the experiment. *Sphingopyxis granuli* strains were grown at 30°C in MML medium or in minimal medium, MM (4) supplemented with tetralin in the vapour phase, 40 mM β-hydroxybutyrate (βHB) or 16 mM sebacic acid as carbon and energy sources. When required, sucrose was added to M63 medium at a final concentration of 5 % and IPTG to 0.0002 %. The media for *E. coli* and *S. granuli* were supplemented with antibiotics when required at the concentrations indicated in Supplementary Material (Table S1).

### Construction of plasmids and strains

Strains, plasmids and oligonucleotides used in this word can be found in Supplementary Material (Table S2). The deletion mutant of SGRAN_2041 (*lysR*) and SGRAN_2210 (*hfq*) were constructed as described in (*33*). Briefly, the upstream and downstream flanking regions of SGRAN_2041 and *hfq* were amplified with specific oligonucleotides (Supplementary Material, Table S3). The resulting upstream fragment was digested with EcoRI and HindIII (for *hfq*) and EcoRI and BamHI (for *lysR*), while the resulting downstream fragment was digested with HindIII and XbaI (for *hfq*) and BamHI and XbaI (for *lysR*). Both fragments were then cloned into the pUC19 plasmid cut by EcoRI and XbaI. After validating the cloning via PCR and subsequent sequencing, the confirmed insert was subcloned into the suicide pMPO1412 vector, utilizing the same restriction sites. The resulting plasmid (pMPO1161 for *hfq* and pMPO1822 for *lysR*) was introduced into competent TFA cells via electroporation. Transformants were selected on agar plates supplemented with 20 mg/L of kanamycin (Km) and then screened for sensitivity to 200 mg/L of streptomycin (Str). Plasmid integration into the genome was validated by PCR amplification of the kanamycin resistance cassette using the oligonucleotides KmFw-pk18 and KmRev-pK18 (Supplementary Material. Table S3). To force the second recombination, the pSW-I vector, which expresses the I-SceI endonuclease, was introduced by electroporation. Basal expression of the endonuclease was sufficient to cut the I-SceI sites, so no induction was required. Candidates that had undergone the second recombination were identified by their loss of kanamycin resistance. The successful gene deletion was initially confirmed by PCR screening of the candidate colonies using specific oligonucleotides (Supplementary Material. Table S3). Finally, the pSW-I vector was cured from the mutant strain by repeated subculturing in the absence of ampicillin. The successful loss of the plasmid was confirmed by the resulting loss of ampicillin resistance. Final validation of the deletion was achieved by sequencing the PCR products.

The strains MPO667 and MPO668 derived from *E. coli* PM1805 were constructed as described in (*32*). Briefly, to construct the translational fusions SGRAN_4018::*lacZ* (MPO667) and SGRAN_3755::*lacZ* (MPO668) in PM1805, the corresponding genes, encoding a TonB-dependent receptor (TBDR) and an acetoacetyl-CoA reductase involved in PHB biosynthesis, *phbB*, respectively, were amplified from the transcription start site (TSS) to the end, excluding the stop codon, using primers PM1805_4018_F and PM1805_4018_R (SGRAN_4018) and PM1805_3755_F and PM1805_3755_R (SGRAN_3755). The primers contained complementary sequences for recombination-mediated replacement of the *cat-sacB* cassette, placing each gene under the P_BAD_ promoter and fused to *lacZ*. The PCR products were electroporated into PM1805, and recombinase expression was induced by incubation at 42 °C with shaking for 15 min. Recombinants were selected on M63 agar supplemented with 5% sucrose and 0.2% glycerol, and correct replacement was verified by PCR and sequencing with oligonucleotides pBAD.for and PM1805_lacZ_Rv.

The plasmid for overexpression of SGRAN_2041 (LysR activator), pMPO1169, was constructed according to the instructions in the IMPACT kit manual (New England Biolabs). The fragment was amplified by PCR using primers SGRAN_2041_ATG_F and SGRAN_2041_BamHI_R, then digested with BamHI and cloned into the expression vector pTYB21, which had been digested with SapI, blunted with Klenow, and then digested again with BamHI.

The plasmid used for the heterologous expression of *suhB* in PM1805-derived strains, pMPO1178, was constructed by amplifying the *suhB* gene using primers AatII_suhB_Fw and EcoRI_suhB_Rv. The resulting PCR fragment was digested with AatII and EcoRI and cloned into plasmid pNM46, previously digested with the same restriction enzymes. For the heterologous expression of *hfq* in PM1805-derived strains, plasmid pMPO1158 was constructed by amplifying the gene from its transcriptional start site (TSS) to the end of its open reading frame (ORF) using primers EcoRI_tsshfq_fw and BamHI_hfq_rv. The insert was digested with EcoRI and BamHI and cloned into pSEVA224, cut with the same enzymes. This construct places *hfq* expression under the control of the P_trc_ promoter. Finally, to complement the *hfq* mutant, the *hfq* gene was amplified by PCR from its promoter to the end of the coding sequence using primers EcoRI_phfq_fw and BamHI_hfq_rv and cloned into pSEVA221. Both the PCR fragment and the vector were digested with EcoRI and BamHI, resulting in plasmid pMPO1179.

### Stress resistance assays

To test the resistance to different stressors, the stress phenotypic assays were done as described in (*23*) with some modifications. Briefly, to evaluate resistance to osmotic and heavy metal stres**s** (copper), late-exponential phase cultures were prepared. These were adjusted to an OD 600 of 1, serially diluted, and 10 μL were dropped onto MML plates containing either 550 mM of NaCl or 3.5 mM of CuSO_4_. The plates were then incubated at 30 °C for 5-6 days until biomass growth was observed. For desiccation assays, 5 μL of the serial dilutions were placed on filters with 0.45 μm pore size (Sartorius Stedim Biotech GmbH, Germany). They were left to air-dry in a laminar flow cabinet for 2 hours (while the control was only dried for 5 minutes) and then kept at room temperature for an additional 20 hours. Afterward, the filters were transferred to plates with MML rich medium supplemented with 0.002% bromophenol blue and incubated at 30 °C for 5-6 days. In the case of oxidative shock resistance assay, late-exponential phase cultures were diluted to an OD 600 nm of 0.1. Once the OD reached 0.5, H_2_O_2_ was added to the medium to a final concentration of 10 mM. Recovery from the shock was assessed by comparing the OD 600 nm of treated cultures after 5 hours of growth to that of untreated cultures, with the result expressed as a percentage. The experiments were performed at least in triplicate, and the most representative examples are shown.

### Proteomic analysis

For label-free quantitative (LFQ) proteomics, TFA wild type (WT) and the Δ*suhB* mutant (MPO642) were cultivated in minimal medium (MM) supplemented with 40 mM 3-hydroxybutyrate (BHB) until reaching exponential (OD600 nm ∼0.8) and stationary phase (OD600 nm ∼3.0). Cells were harvested by centrifugation at 10,000 × g for 10 min at 4 °C, washed once with 50 mM Tris-HCl (pH 8.0), and cell pellets were stored at −80 °C until further processing. For protein extraction, frozen pellets were resuspended in lysis buffer (100 mM Tris-HCl, pH 7.9, 150 mM NaCl and 1.5% SDS), supplemented with protease inhibitors (Merck) and disrupted by sonication (pulse mode, 2 seconds on/10 seconds off, 4 min total, 10% amplitude). Lysates were clarified by centrifugation at 10,000 × g for 30 min at 4 °C, and supernatants were stored at −80 °C. Protein concentration was determined using the RC DC protein assay (Bio-Rad), and quantification of the total protein was verified by loading 10 μg of each sample in an SDS-PAGE gel (12% acrylamide).

After quantification, protein pellets were resuspended in 6M urea, 50 mM ammonium bicarbonate (AB). Disulfide bonds were reduced by adding 10 mM DTT (in 50 mM AB) to the protein solution and incubating for 60 minutes at room temperature. For carbamidomethylation of cysteine –SH groups, 30 mM IAA (in 50 mM AB) was added to the samples, which were then incubated in the dark for 30 minutes. Samples were digested overnight at 37 °C using bovine trypsin (Sequencing Grade Modified Trypsin, Promega) at a 1:12 enzyme-substrate ratio. The reaction was stopped by adding formic acid to a final concentration of 0.5%. OMIX C18 tips (Agilent Technologies) were used for concentrating and desalting the peptide extracts. Samples were dried, diluted in 0.1% formic acid, and injected into a nano-HPLC system. Protein digests were separated on a Thermo Scientific™ Easy nLC system using a 50 cm C18 Thermo Scientific™ EASY-Spray™ column. The following solvents were employed as mobile phases: Water with 0.1% formic acid (phase A) and acetonitrile with 20% H2O and 0.1% formic acid (phase B). Separation was achieved with an acetonitrile gradient from 10% to 35% over 240 minutes, from 35% to 100% over 1 minute, and 100% B over 5 minutes, at a flow rate of 200 nL/min. A Thermo Scientific™ Q Exactive™ Plus Orbitrap™ mass spectrometer was used to acquire the top 10 MS/MS spectra in DDA mode. LC-MS data were analyzed using FragPipe v23 and FragPipe-Analyst (http://fragpipe-analyst.nesvilab.org/), with the parameters recommended for LFQ (*34*) and using the proteome of TFA as database (UniProt Proteome: <u>UP000058599</u>). Proteins were considered as upregulated if the Log2 fold change (Δ*suhB*/WT) was greater than 1, with an adjusted p-value below 0.05. Conversely, proteins were considered downregulated if the Log2 fold change (Δ*suhB*/WT) was less than −1, with an adjusted p-value below 0.05.

### Affinity chromatography assay targeting the *suhB* promoter region

Proteins interacting with the *suhB* promoter were identified using a biotinylated DNA immobilization assay with magnetic beads. The DNA fragment containing *suhB* promoter was amplified by PCR with primers SuhB_fw and SuhB_rvBtn, and reactions were pooled to obtain 15 μg of DNA per assay. Binding of the biotinylated fragment to Dynabeads M-280 (Invitrogen) was done following the manufacturer’s instructions. For the assays, crude extracts were prepared from WT cultures grown in MM supplemented with either 16 mM sebacic acid (inducing condition) or tetralin (non-inducing condition) until reaching exponential phase (OD600 nm ∼0.8). Then, the protocol described by (*35*) was followed with minor modifications. Proteins were extracted by cell disruption through sonication (2 seconds on/10 seconds off pulse cycle) in binding buffer, followed by centrifugation for 20 min at 4 °C. After binding to the DNA-coated magnetic beads, proteins were eluted by adding 25 μL of loading buffer and heating from 50 °C to 95 °C, maintaining the samples at 95 °C for 5 min. Samples were then centrifuged at maximum speed for 5 min at 4 °C. Eluted proteins were analyzed by SDS-PAGE on a 10% polyacrylamide gel and differential bands between extracts were excised from the gels and submitted for identification by MALDI-TOF/TOF mass spectrometry at the Proteomics Service of Universidad Pablo de Olavide.

### Overexpression and purification of the LysR transcriptional activator SGRAN_2041

The LysR-type transcriptional regulator SGRAN_2041 was purified using the IMPACT system (New England Biolabs) according to the manufacturer’s guidelines, with minor adjustments. For protein overproduction, plasmid pMPO1169 was introduced into *E. coli* strain ER2566. A saturated overnight culture was diluted to an initial OD600 of 0.1 in 400 mL of LB medium and incubated at 37 °C with shaking until reaching an OD600 of about 0.5. At this point, cultures were cooled on ice for 30 min and subsequently induced with 0.1 mM IPTG, followed by incubation at 16 °C for 22 h under agitation. Cells were collected by centrifugation, and induction of protein expression was verified by SDS-PAGE. Cell pellets were resuspended in purification buffer (100 mM Tris-HCl pH 8, 0.5 M NaCl, 10% glycerol) and disrupted by sonication. After centrifugation to clarify the lysates, the supernatant was loaded by gravity onto a column containing 5 mL of chitin resin pre-equilibrated with 25 mL of purification buffer. The column was washed with the same buffer before initiating the on-column cleavage. To release the target protein, the resin was incubated at 4 °C with cleavage buffer (purification buffer supplemented with 50 mM DTT) for approximately 72 h. To improve cleavage efficiency, a second incubation step was performed with fresh buffer containing 100 mM DTT for 48 h at room temperature. The protein was then eluted, and its enrichment was confirmed by SDS-PAGE. For downstream assays, dialysis was not required since the purification buffer closely resembled the assay buffer, differing only in salt concentration. Therefore, fractions 1-4 were pooled and diluted with working buffer (100 mM Tris-HCl pH 8, 0.05 M NaCl, 10% glycerol) to a final volume of 40 mL. Finally, the preparation was concentrated using Amicon Ultra centrifugal filters with a 10 kDa cutoff (Millipore). Then, the protein was quantified using the RC DC Protein Assay kit (BioRad) and aliquots were stored at −80 °C.

### Electrophoretic mobility shift assay

The binding of the LysR transcriptional activator to the SuhB promoter was evaluated by fluorescence electrophoretic mobility shift assay (fEMSA) as described in (*36*). To prepare fluorescent probes for fEMSA, a WT probe containing the defined LysR binding site and a mutant probe with targeted mutations were designed. The probes were generated by hybridizing a long (51-base) oligonucleotide (either WT or mutant) with a shorter complementary oligonucleotide containing a 5’ 6-FAM fluorescent dye label (Supplementary Material. Table S3). Oligonucleotides were resuspended in TE buffer to a final concentration of 100 µM. For hybridization, a mixture containing 0.6 µL of the short fluorescent oligo, 1.2 µL of the long oligo, and 28.2 µL of STE buffer (10 mM Tris-HCl pH 8, 1 mM EDTA, 100 mM NaCl) was incubated in a thermocycler. The cycle involved heating to 95 °C for 2 minutes, followed by a gradual ramp down to 25 °C, and a final hold at 4 °C. The hybridized oligos were then filled by Klenow polymerase (New England Biolab) to complete the double-stranded probe. The reaction was incubated at 37 °C for 60 minutes, then inactivated by adding 3.4 µL of 0.5 M EDTA and incubating at 75 °C for 20 minutes. The completed probes were diluted in STE to a final concentration of 0.1 µM, aliquoted, and stored at −20 °C in the dark.

Binding reactions were prepared in a 20 µL volume containing 5 nM of the fluorescent probe (WT or mutant), 5X Binding Buffer (*36*), poly-dI-dC as competitor, and increasing amounts of purified LysR protein. The mixtures were incubated at room temperature for 30 minutes in the dark. Samples were then loaded in a 5% native polyacrylamide gel after adding a TE plus 50% glycerol loading buffer. The gel was pre-run for 1 hour at 80 V and then run for approximately 40 minutes at the same voltage. Gels were imaged directly in their glass plates using a Typhoon scanner with the Cy2 fluorescence setting.

### Measurement of β-Galactosidase activity

TFA cultures were grown at 30°C in MM plus 40 mM of βHB, aliquots were collected at different points of the growth curve and β-galactosidase activity was determined (*37*). SuhB induction was measured in cultures that had been inoculated with cells grown on tetralin. For the *E. coli* PM1805-derived strains, saturated overnight cultures were grown at 37°C in LB medium containing 0.0002% arabinose and 0.1 mM IPTG and then collected to assess β-galactosidase activity. All graphs were generated using the average of three independent biological replicates.

### Transmission electron microscopy (TEM)

Cultures of TFA WT, MPO642 (Δ*suhB*), and MPO209 (*miniTn5::phaC*) were grown in minimal medium supplemented with 40 mM β-hydroxybutyrate and the appropriate antibiotics. Cultures were inoculated from overnight pre-cultures to an initial optical density at 600 nm (OD_600_) of 0.05. After 16 h (exponential phase) and 36 h (stationary phase) of incubation at 30 °C with shaking, cells were harvested by centrifugation and resuspended in 1 mL of 2.5% glutaraldehyde. Samples were fixed for 2 h at 4 °C, centrifuged, and the supernatant was discarded. The pellets were then washed with 0.1 M cacodylate buffer by resuspension and incubation for 15 min at room temperature, followed by centrifugation and supernatant removal. This washing step was repeated three times. Then, the samples were embedded in agarose and post-fixed in 1% osmium tetroxide for 1 hour, followed by three washes in 0.1 M cacodylate buffer. Subsequently, they were incubated in 2% uranyl acetate for 2 hours. The samples were then dehydrated through a graded acetone series up to 100% acetone and embedded in Spurr resin, which was polymerized for 7 hours at 70°C. Ultrathin sections (70 nm) were obtained using a Leica UC7 ultramicrotome and examined with a Zeiss Libra 120 transmission electron microscope operated at 80 kV. PHB granule quantification was performed by measuring the area occupied by granules relative to the total cellular area in six TEM images from WT and six from the Δ*suhB* mutant, using FIJI-ImageJ software according to the protocol described in (*38*).

### Bioinformatic analyses for target prediction and GO terms enrichment

To predict possible SuhB targets, we followed a combined bioinformatic approach. The tool ANNOgesic (*39*), which integrates IntaRNA (*40*), RNAplex (*41*), and RNAup (*42, 43*), was used to identify potential novel targets. Additionally, CopraRNA (*40*) was employed with default parameters (predicting interactions on a sequence window comprising 200 bp upstream of the start codon and 100 bp downstream), using as queries SuhB homologs from TFA (NZ_CP012199), *Sphingopyxis* sp. 113P3 (NZ_CP009452), *S. macrogoltabida* (NZ_CP012700), *Sphingopyxis* sp. QXT-31 (NZ_CP019449), *Sphingopyxis* sp. MG (NZ_CP026381), and *S. alaskensis* RB2256 (NC_008048). Only targets showing differential expression in Δ*suhB* versus WT in proteomic data (both exponential and/or stationary phases) were considered. Furthermore, Hfq_3xFlag co-immunoprecipitation data (*31*) were analyzed to evaluate whether the predicted targets could bind Hfq, as observed for SuhB (*31*).

Proteins upregulated and downregulated in Δ*suhB* were subjected to biological enrichment analysis based on their annotated GO terms. The analysis was performed using Fisher’s exact test with the TopGO R package (version 2.56.0) applying the classic algorithm (*44*), and the results were visualized using ggplot2 (*45*).

## Results

### SuhB is essential for survival in different environmental stresses

The only small RNA characterized so far in TFA is SuhB, which is known to participate in the carbon catabolite repression of tetralin degrading genes by repressing the translation of their transcriptional activator ThnR (*31*). However, no other phenotypes have been described for SuhB in this bacterium. To further investigate its role, we tested whether the absence of SuhB affects the ability of TFA to resist different stresses previously characterized in TFA by comparing the stress resistance of the Δ*suhB* mutant (MPO642) with that of the wild-type (WT) strain. As a control, we introduced in our experiments the double mutant lacking the sigma factors of the GSR (Δ*ecfG1*Δ*ecfG2*, MPO860). This GSR regulatory mutant does not activate the GSR and have a survival defect in stress conditions (*23*). We observed that Δ*suhB* displayed increased sensitivity to heavy metal (copper), desiccation, osmotic stress, and oxidative stress when compared to the WT in a similar way than Δ*ecfG1*Δ*ecfG2* (Figure 1). Complementation of Δ*suhB* restored WT stress resistance (Supplementary Material. Figure S1), confirming the regulatory role of SuhB in stress survival. SuhB, as many others sRNAs, is associated with the RNA chaperone Hfq. Hfq is involved in regulating multiple functions in the cells and it has been previously described to be involved in stress tolerance (*46*). Therefore, we also assessed the stress resistance of an *hfq* deletion mutant (MPO646). In TFA, deletion of *hfq* does not affect growth under non-stress conditions but shows a sensitivity profile to all tested stresses similar to that of Δ*ecfG1*Δ*ecfG2* (Figure 1). Moreover, complementation with *hfq* restores stress resistance in the deletion mutant (Supplementary Material, Figure S2). The stress sensitivity of the *hfq* mutant likely reflects its broad role in RNA regulation, which includes facilitating interactions between SuhB and its targets.

**Figure 1:**
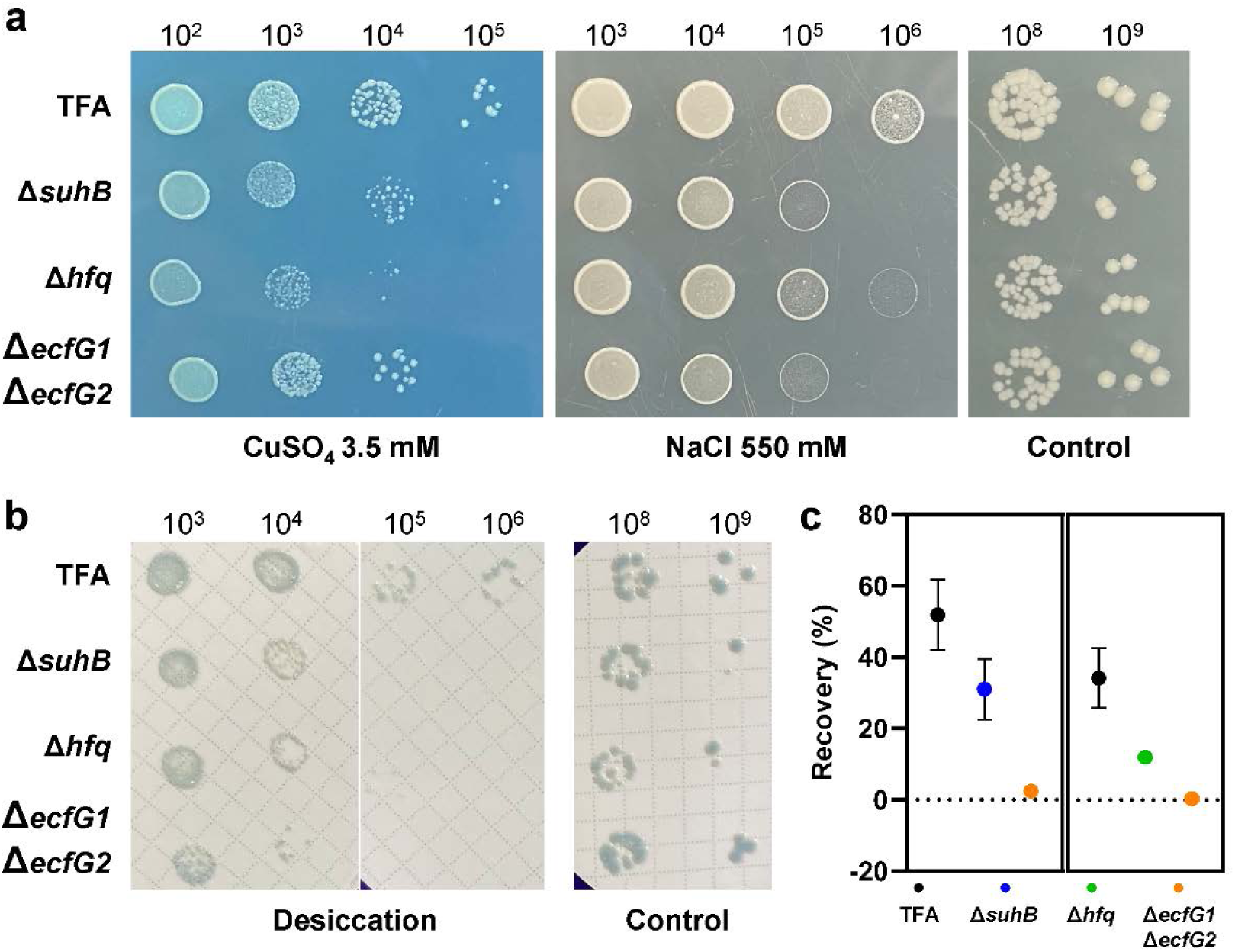
Phenotypic stress assays comparing *suhB* and *hfq* deletions mutants with the wild type TFA and the mutant lacking the GSR sigma factors. (a) Stress sensitivity was assessed by spotting serial dilutions of the different strains onto MML rich agar supplemented with 3.5 mM CuSO_4_ (heavy metal stress) or 500 mM NaCl (osmotic stress). (b) Desiccation tolerance was evaluated by placing serial dilutions on filters and allowing them to air dry for 24 h. (c) The ability to recover from oxidative stress was determined by comparing treated cultures to untreated controls 5 h after exposure to 10 mM H_2_O_2_.

### The absence of SuhB significantly alters the proteome of TFA

To gain deeper insight into the regulatory role of SuhB, we conducted a label-free quantitative (LFQ) proteomic analysis comparing WT and Δ*suhB* strains during both exponential and stationary phases of growth in minimal medium (MM) supplemented with 40 mM β-hydroxybutyrate (BHB) as the sole carbon source, a condition of induction of *suhB* expression (*31*). Quality control of the data was done by Principal Component Analysis (PCA), using the 500 most variable proteins to assess sample clustering according to the experimental condition. In both exponential (Figure 2a) and stationary phases (Figure 2b), the first principal component (PC1) clearly separated the WT and Δ*suhB* samples, accounting for 46.7% and 54.4% of the total variance, respectively. The second principal component (PC2) contributed 17.7% of the variance in both phases and primarily captured replicate variability within each group. This separation confirms a consistent and robust effect of the *suhB* deletion on the global proteomic profile in both growth phases.

**Figure 2:**
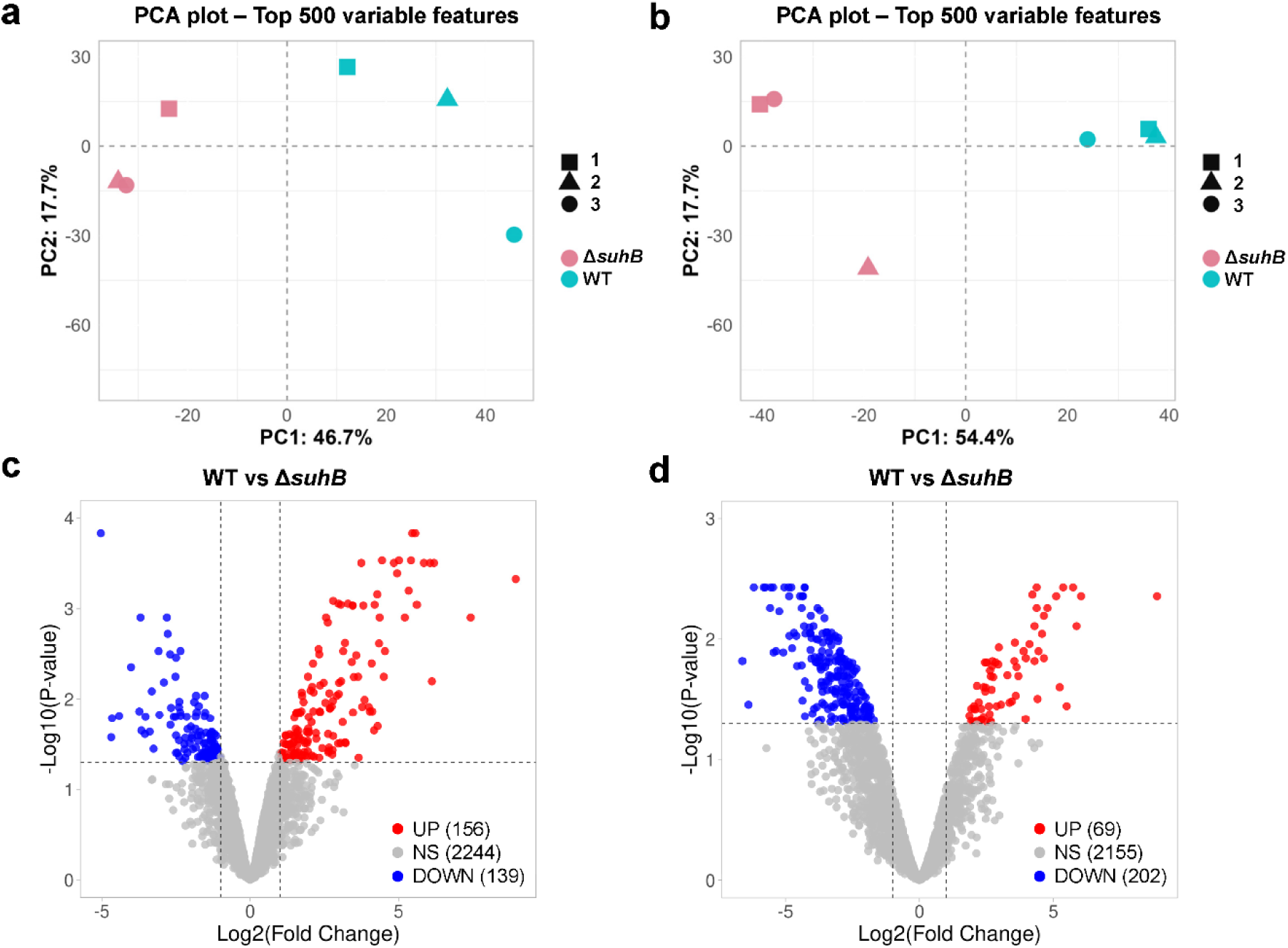
Principal Component Analysis (PCA) and differential gene expression analysis of wild type TFA and Δ*suhB* strains. (a) Principal component analysis (PCA) of proteomic data from the exponential phase. (b) PCA of proteomic data from the stationary phase. (c) Volcano plot showing upregulated (red), downregulated (blue), and non-significantly changed (grey) proteins in Δ*suhB* during the exponential phase. (d) Volcano plot showing upregulated (red), downregulated (blue), and non-significantly changed (grey) proteins in Δ*suhB* during the stationary phase.

In the exponential phase, a total of 2,539 proteins were identified out of the 4,190 annotated in the TFA genome, representing approximately 61% coverage. Among these, 156 proteins were significantly upregulated (Log2FC >1, adjusted p-value < 0.05) and 139 downregulated (Log2FC <-1, adjusted p-value < 0.05) in Δ*suhB* compared to the WT (Figure 2c). During the stationary phase, 2,426 proteins were identified (58% coverage), with 69 proteins upregulated and 202 downregulated in the mutant (Figure 2d). These differentially expressed proteins constitutes the SuhB regulon, including directly regulated targets as well as proteins whose abundance is indirectly affected by SuhB regulation (Supplementary Data 1).

To further explore the functional implications of these proteomic changes, we performed Gene Ontology (GO) enrichment analysis. In the exponential phase, proteins upregulated in the Δ*suhB* strain were significantly enriched in cellular component terms associated with the membrane, and in molecular functions related to fatty acid-CoA ligase activity. Conversely, downregulated proteins were enriched in biological processes related to small molecule metabolism, as well as intracellular proteins and hydrolase activity based on cellular component and molecular function enrichments, respectively (Supplementary Material. Figure S3 and Figure S4).

In the stationary phase, the upregulated proteins were also enriched in terms related to the membrane, such as transmembrane transporters. Moreover, the downregulated proteins were once again enriched in biological processes related to small molecule metabolism, as well as in hydrolase activity according to the molecular function category (Supplementary Material. Figure S5 and Figure S6).

Remarkably, among the differentially expressed proteins, a high number of TBDRs are upregulated in the Δ*suhB* mutant, specially under exponential growth conditions. TFA contains 87 TBDRs annotated in the genome, which is an elevated number also found in other oligotrophic bacteria (*47*). Out of the 66 TBDRs detected under this condition, 29 are upregulated, none is downregulated, and 37 remains equal. In stationary phase, 2 TBDRs are upregulated, 6 are downregulated, and 60 does not change its expression within the 68 that were detected. (Supplementary Data 2).

All together, these results suggest that SuhB play an important role in adjusting the membrane proteome in TFA, likely contributing to the physiological adaptation under specific growth conditions.

### Potential mRNA targets of SuhB in TFA

Until now, the only experimentally validated target of SuhB in TFA was the mRNA of the transcriptional activator of the tetralin degradation pathway ThnR. In this study, we combined *in silico* prediction tools with quantitative proteomic data to identify potential additional targets regulated by SuhB. Specifically, we used COPRA and ANNOgesic to predict potential SuhB targets based on sequence complementarity and conservation of the sRNA-target pair in other *Sphingopyxis* species (Supplementary Data 3 and 4). Only those predicted targets that also showed significant differential protein abundance between the Δ*suhB* mutant and the WT strain were selected (Table 1). The predicted SuhB targets were grouped into 8 functional categories which are metabolism of cofactors and vitamins, transcription/translation, transcriptional regulators, fatty acid metabolism, PHB biosynthesis, transporters/membrane proteins, organic compounds metabolism and other functions (Figure 3). Among these candidates, *phbB* (SGRAN_3755), encoding an acetoacetyl-CoA reductase involved in PHB biosynthesis, was of particular interest. PhbB catalyzes the reduction of acetoacetyl-CoA to (R)-3-hydroxybutyryl-CoA and accumulated in both exponential and stationary phases in Δ*suhB*, suggesting repression by SuhB, probably by blocking its Shine-Dalgarno sequence as predicted by COPRA (Figure 4a). Several TBDRs showing altered expression in the mutant were also predicted SuhB targets (Table 1).

**Figure 3:**
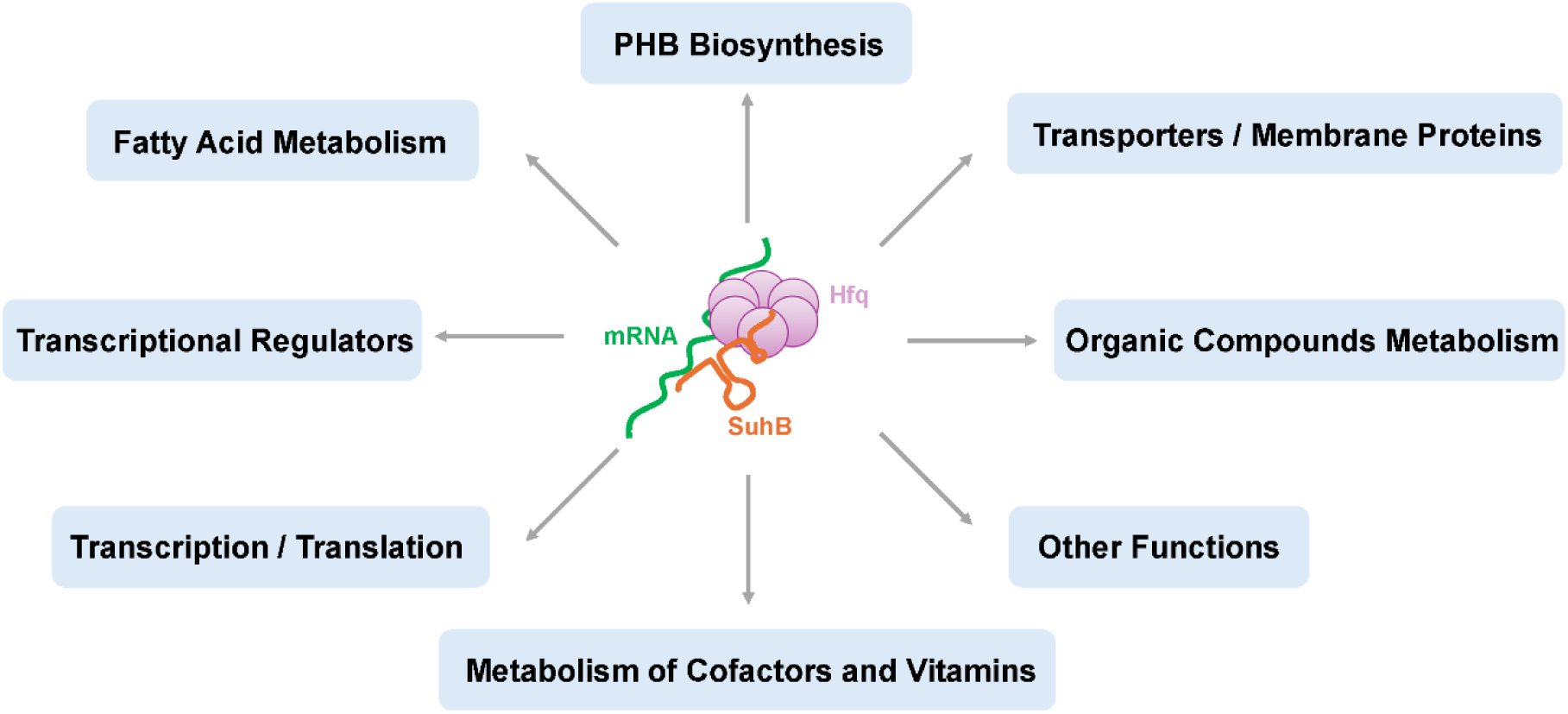
Functional classification of predicted SuhB targets. Classification based on biological functions of the X possible targets for SuhB, predicted based on COPRA and ANNOgesic analysis and differential expression in proteomic data on both exponential and stationary phase.

**Figure 4:**
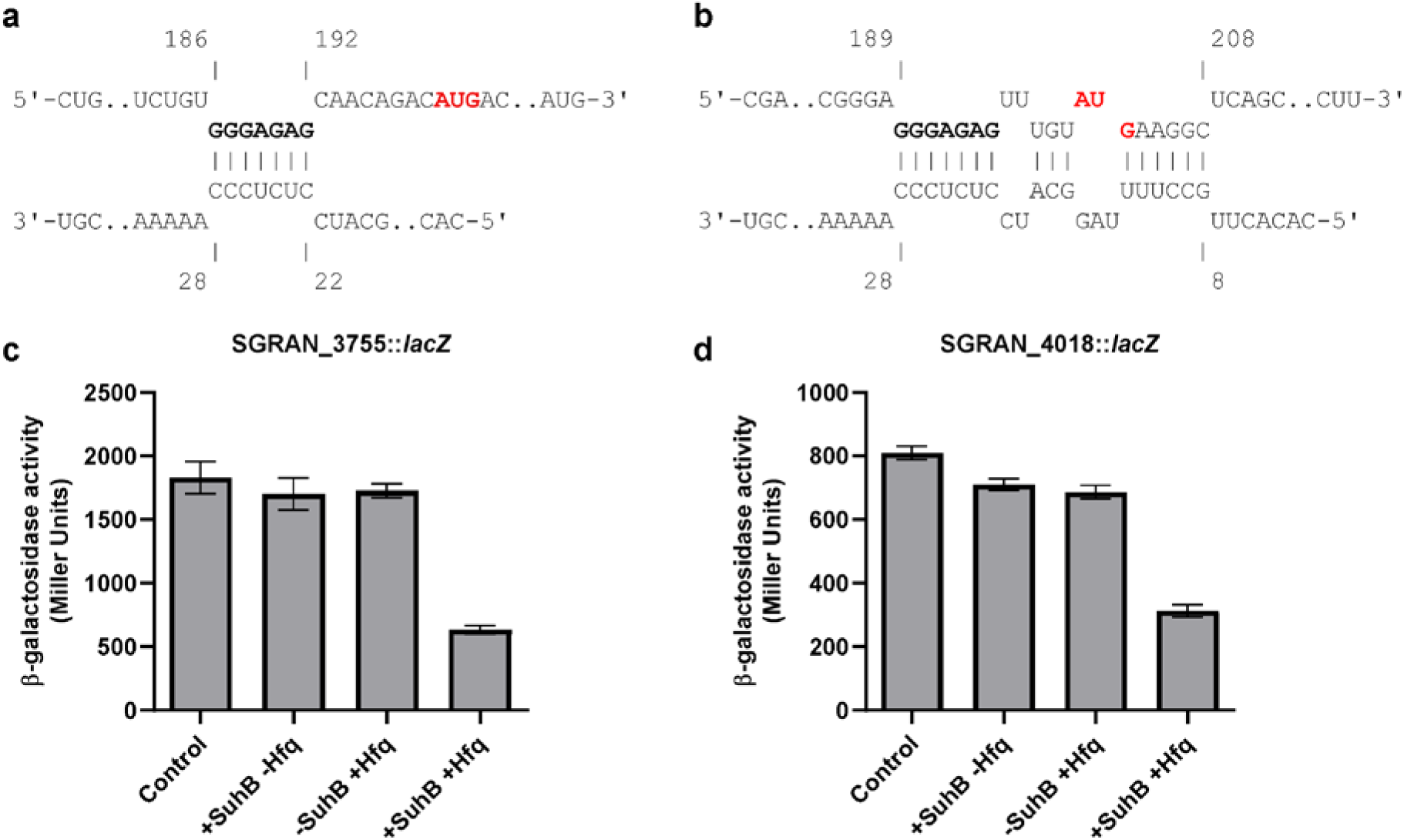
Repression of P_BAD_-*SGRAN_4018*_*lacZ* and P_BAD_-*SGRAN_3755*_*lacZ* expression by SuhB in the absence or presence of the TFA Hfq. Interaction regions of (a) SGRAN_3755 mRNA and (b) SGRAN_4018 mRNA with SuhB. The Shine-Dalgarno sequences of the mRNAs are marked in bold, and the start codons are highlighted in red. mRNA is represented from 5’ to 3’, and SuhB from 3’ to 5’. β-galactosidase activity assays were performed to measure the expression of the translational fusions in MPO667 (P_BAD_-*SGRAN_4018*_*lacZ*) and MPO668 (P_BAD_-*SGRAN_3755*_*lacZ*) upon induction of SuhB and Hfq. Bacterial cultures were grown overnight, and the expression of SuhB and Hfq was induced with 0.1 mM IPTG, while the expression of the translational fusions was induced with 0.0002% L-arabinose. Measurements were performed in three biological replicates and two independent experiments.

**Table 1:** Predicted SuhB targets based on COPRA and ANNOgesic analysis (at least one of them), differential expression in proteomic data on both exponential and stationary phase and binding to Hfq according to Hfq CoIP.

| Gene ID | Description | Log2(FC) | Functional Category | COPRA | Annogesic | Hfq |
| --- | --- | --- | --- | --- | --- | --- |
| **Upregulated in ΔsuhB in exponential phase** |  |  |  |  |  |  |
| SGRAN_3397 | Diacylglycerol O-acyltransferase | 6.12 | Fatty acid metabolism | Yes | No | Yes |
| SGRAN_3257 | AMP-dependent synthetase and ligase | 5.85 | Fatty acid metabolism | No | Yes | Yes |
| SGRAN_1051 | TonB-dependent receptor-like protein | 5.62 | Transporters / Membrane proteins | No | Yes | Yes |
| SGRAN_1608 | TonB-dependent receptor | 5.56 | Transporters / Membrane proteins | Yes | Yes | Yes |
| SGRAN_3454 | Alcohol dehydrogenase (azurin) | 5.22 | Organic compounds metabolism | Yes | No | Yes |
| SGRAN_3880 | TonB-dependent receptor | 5.02 | Transporters / Membrane proteins | Yes | Yes | No |
| SGRAN_4018 | TonB-dependent receptor | 4.55 | Transporters / Membrane proteins | Yes | No | Yes |
| SGRAN_1394 | Uncharacterized protein | 4.3 | Other functions | Yes | No | Yes |
| SGRAN_0137 | TonB-dependent receptor, plug | 4 | Transporters / Membrane proteins | Yes | Yes | Yes |
| SGRAN_2788 | TonB-dependent receptor-like protein | 3.46 | Transporters / Membrane proteins | No | Yes | Yes |
| SGRAN_2011 | Major facilitator transporter | 3.43 | Transporters / Membrane proteins | Yes | Yes | Yes |
| SGRAN_1428 | Uncharacterized protein | 3.2 | Other functions | No | Yes | Yes |
| SGRAN_3990 | Major facilitator superfamily MFS-1 | 3 | Transporters / Membrane proteins | Yes | No | Yes |
| SGRAN_3755 | 3-oxoacyl-[acyl-carrier-protein] reductase FabG | 2.8 | PHB biosynthesis | Yes | No | Yes |
| SGRAN_2804 | LysR-type activator (ThnR) | 2.79 | Transcriptional regulators | No | Yes | Yes |
| SGRAN_1424 | Acyl-CoA transferase/carnitine dehydratase | 2.77 | Fatty acid metabolism | No | Yes | Yes |
| SGRAN_1300 | Haloalkane dehalogenase | 2.62 | Organic compounds metabolism | Yes | Yes | Yes |
| SGRAN_1740 | TonB-dependent receptor | 2.56 | Transporters / Membrane proteins | Yes | No | Yes |
| SGRAN_0324 | Polyketide cyclase | 2.36 | Other functions | Yes | No | Yes |
| SGRAN_1972 | Membrane protein | 2.36 | Transporters / Membrane proteins | Yes | No | No |
| SGRAN_2616 | Uncharacterized protein | 2.22 | Other functions | Yes | Yes | Yes |
| SGRAN_3051 | Eicosanoid-glutathione metabolism membrane protein | 2.07 | Transporters / Membrane proteins | Yes | No | Yes |
| SGRAN_1960 | Long-chain fatty acid--CoA ligase | 2.04 | Fatty acid metabolism | Yes | No | Yes |
| SGRAN_3419 | Acyl-CoA dehydrogenase | 1.83 | Fatty acid metabolism | Yes | Yes | Yes |
| SGRAN_1605 | 3-oxocholest-4-en-26-oate--CoA ligase | 1.74 | Fatty acid metabolism | Yes | No | Yes |
| SGRAN_0267 | Glutamate synthase | 1.68 | Organic compounds metabolism | Yes | No | Yes |
| SGRAN_4001 | Alkyl sulfatase | 1.67 | Organic compounds metabolism | Yes | No | Yes |
| SGRAN_3502 | TetR family transcriptional regulator | 1.63 | Transcriptional regulators | No | Yes | Yes |
| SGRAN_0508 | Acyl-CoA dehydrogenase domain-containing protein | 1.56 | Fatty acid metabolism | No | Yes | Yes |
| SGRAN_2191 | Glycosyl transferase, group 1 | 1.39 | Other functions | Yes | Yes | Yes |
| SGRAN_1989 | Acyl-PO4 G3P acyltransferase | 1.09 | Fatty acid metabolism | Yes | No | No |
| SGRAN_2302 | Uncharacterized protein | 1.09 | Other functions | No | Yes | Yes |
| **Downregulated in ΔsuhB in exponential phase** |  |  |  |  |  |  |
| SGRAN_1462 | Amidohydrolase | -4.65 | Other functions | No | Yes | Yes |
| SGRAN_2615 | Aspartyl/glutamyl-tRNA amidotransferase subunit C | -4.41 | Other functions | Yes | No | Yes |
| SGRAN_0418 | Sulfatase | -1.83 | Other functions | Yes | No | Yes |
| SGRAN_2008 | Biotin carboxylase | -1.7 | Metabolism cofactors / vitamins | No | Yes | No |
| SGRAN_0375 | 6,7-dimethyl-8-ribityllumazine synthase | -1.16 | Metabolism cofactors / vitamins | No | Yes | Yes |
| SGRAN_3986 | Glutathione S-transferase | -1.15 | Other functions | Yes | No | No |
| **Upregulated in ΔsuhB in stationary phase** |  |  |  |  |  |  |
| SGRAN_3257 | AMP-dependent synthetase and ligase | 2.67 | Fatty acid metabolism | No | Yes | Yes |
| SGRAN_3755 | 3-oxoacyl-[acyl-carrier-protein] reductase FabG | 2.05 | PHB biosynthesis | Yes | No | Yes |
| **Downregulated in ΔsuhB in stationary phase** |  |  |  |  |  |  |
| SGRAN_3769 | Carboxymethylenebutenolidase | -6.19 | Organic compounds metabolism | Yes | Yes | Yes |
| SGRAN_3348 | Gluconolactonase | -4.79 | Organic compounds metabolism | Yes | No | Yes |
| SGRAN_2761 | Ribosome-recycling factor | -4.11 | Transcription / Translation | Yes | No | Yes |
| SGRAN_2163 | Thiamine-phosphate synthase | -4.07 | Metabolism cofactors / vitamins | Yes | No | No |
| SGRAN_2291 | Probable DNA helicase II like protein | -3.62 | Transcription / Translation | No | Yes | No |
| SGRAN_2086 | Short chain dehydrogenase | -2.89 | Fatty acid metabolism | Yes | No | Yes |
| SGRAN_4060 | Acyl-CoA dehydrogenase-like protein | -2.7 | Fatty acid metabolism | Yes | No | Yes |
| SGRAN_2164 | Oligopeptide transporter, OPT family | -2.42 | Transporters / Membrane proteins | Yes | No | Yes |
| SGRAN_2955 | CheW protein | -2.01 | Other functions | Yes | No | Yes |

To experimentally determine which of the predicted targets were regulated by SuhB, translational *lacZ* fusions were constructed in *E. coli* PM1805. Each construct included the predicted SuhB binding region, typically located around the start codon of the mRNA (Figure 4a, b and Supplementary Figure S7). We selected *phbB* and the TBDRs SGRAN_1608 and SGRAN_4018 for experimental validation. The SGRAN_1608::*lacZ* fusion did not yield sufficient β-galactosidase activity, maybe due to a suboptimal Shine-Dalgarno sequence for *E. coli*, and could not be evaluated (data not shown). In contrast, both *phbB* and SGRAN_4018::*lacZ* fusions produced measurable levels of β-galactosidase, indicating proper expression in *E. coli*. Expression of SuhB or Hfq alone caused only a modest reduction in reporter activity, particularly for SGRAN_4018::*lacZ*, whereas co-expression of SuhB with the TFA Hfq chaperone resulted in a marked decrease (Figure 4c, d). These results confirm both genes as novel SuhB targets and indicate that the heterologous system is fully functional only when the Hfq protein from TFA is present, suggesting that the *E. coli* Hfq cannot efficiently support the interaction between SuhB and its target mRNAs or, alternatively, that higher levels of Hfq are required for proper regulation.

Because PhbB is a key enzyme directing acetoacetyl-CoA into PHB biosynthesis, we examined whether SuhB influences PHB accumulation. Transmission electron microscopy revealed an increase in PHB granules in Δ*suhB* during exponential growth compared with the WT strain (Figure 5). As expected, no granules were observed in the Δ*phaC* negative control (*48*). In stationary phase, neither strain accumulated PHB, consistent with carbon limitation and the consequent mobilization of stored PHB (Supplementary Figure S8). These observations support the conclusion that SuhB represses *phbB* and thereby contributes to controlling PHB biosynthesis.

**Figure 5:**
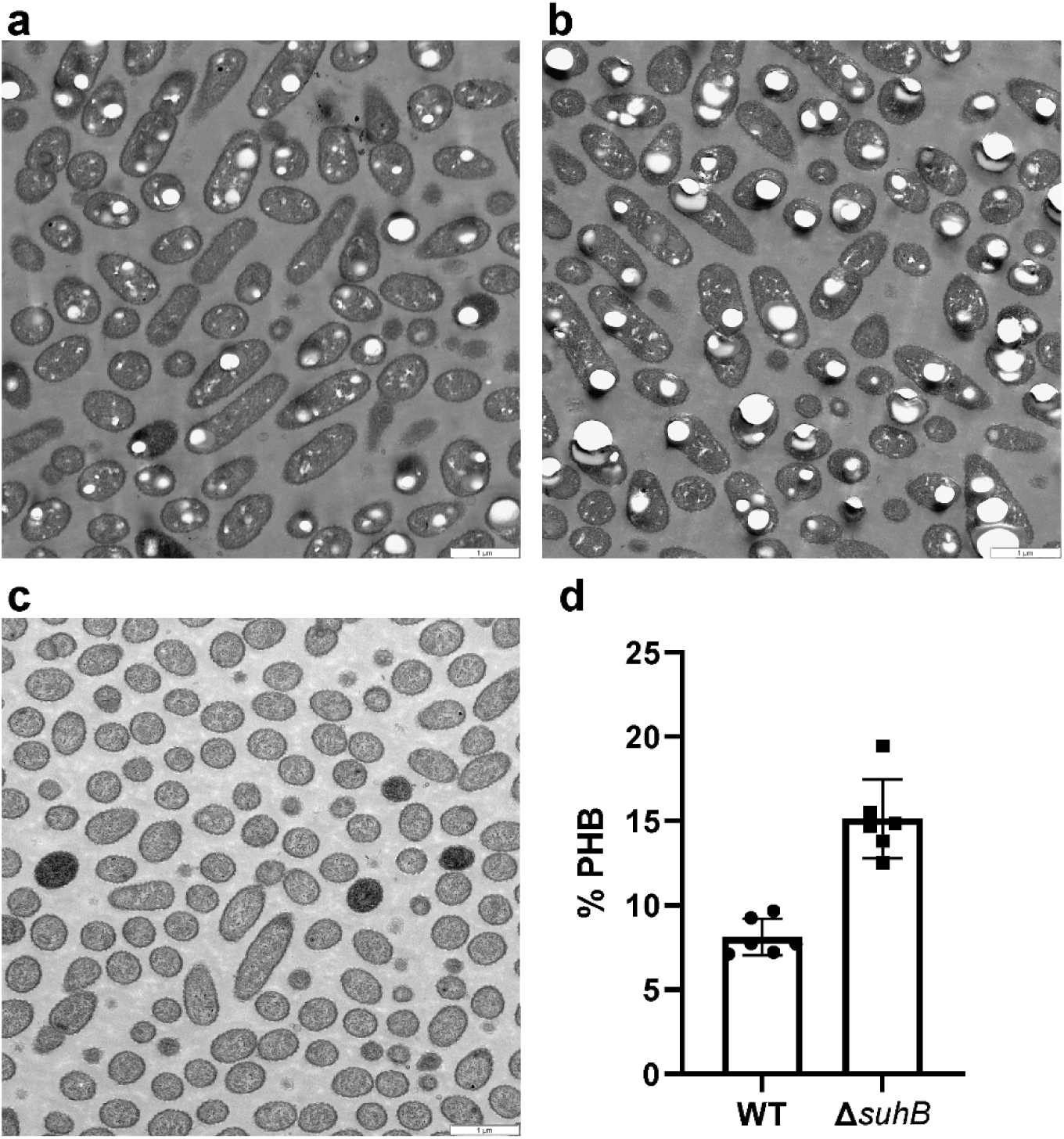
Transmission electron micrographs of cells in exponential phase. (a) *Sphingopyxis granuli* TFA, WT; (b) Δ*suhB* mutant; (c) *miniTn5::phaC* mutant. (d) Percentage of PHB relative to the total cell area, quantified in six independent images using FIJI-ImageJ software. Scale bars represent 1 μm.

### The SuhB regulon is largely independent of GSR controlled genes

The complete SuhB regulon was defined above as the set of proteins whose abundance changes significantly in the Δ*suhB* mutant. Because Δ*suhB* exhibits reduced stress tolerance, we examined whether SuhB could influence genes associated with the general stress response (GSR). Proteomic analyses performed during exponential and stationary phases (the latter being a condition that activates the GSR) showed no differences in the abundance of the GSR regulatory proteins EcfG1, EcfG2, NepR1, NepR2, PhyR1, or PhyR2 between the WT and Δ*suhB* strains (Supplementary Table S4). Consistently, expression of a chromosomal *nepR2::lacZ* translational fusion, commonly used as a reporter for GSR activation, also showed no differences between the two strains (Supplementary Fig. S9).

To further assess possible convergence between SuhB and the GSR, we compared the proteins with altered abundance in Δ*suhB* (in stationary phase) to the GSR regulon defined in stationary phase (Figure 6), using a Venn diagram generated with Venny 2.1 (49). Among the genes upregulated in the GSR regulon, four were also upregulated in Δ*suhB*, including those encoding a secretion system protein (SGRAN_1208), an uncharacterized protein (SGRAN_1676), a TonB-dependent receptor (SGRAN_2777), and the flagellar motor switch protein FliN (SGRAN_4099), whereas nine were downregulated. Conversely, within the genes downregulated in the GSR regulon, four were upregulated and twelve downregulated in Δ*suhB*. Two of the twelve genes downregulated in both datasets encode proteins previously associated with stress tolerance, CspA (SGRAN_2505) and SodB (SGRAN_1121), neither of which was predicted as a direct SuhB target. The remaining ten include several uncharacterized proteins (SGRAN_1892, SGRAN_0758, SGRAN_0034, SGRAN_2412), structural or enzymatic proteins such as the membrane protein SGRAN_1264, the 6-phosphogluconate dehydratase Edd (SGRAN_3544), the glyoxalase family protein SGRAN_3705, the transglycosylase MltA3 (SGRAN_1261), a 17 kDa surface antigen (SGRAN_3922), and a protein belonging to the “required for attachment to host cells” family (SGRAN_1934). Beyond these overlaps, 250 genes were uniquely upregulated and 173 uniquely downregulated in the GSR regulon, while 61 and 181 were exclusively up or downregulated, respectively, in Δ*suhB*.

**Figure 6:**
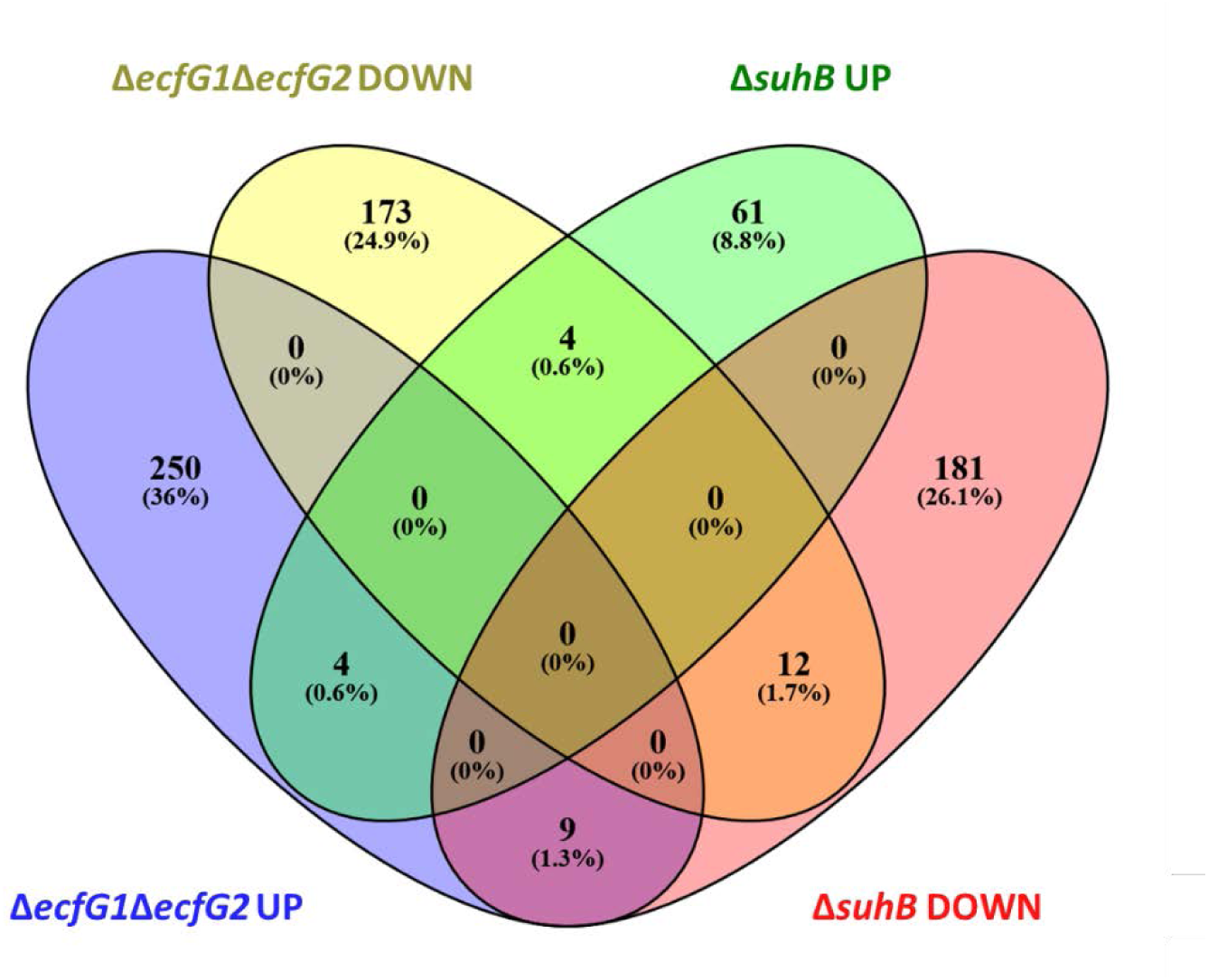
Venn diagram showing the overlap between SuhB and GSR regulons. Proteins differentially expressed in the Δ*suhB* mutant and in the Δ*ecfG1*Δ*ecfG2* mutant, which is deficient in GSR activation. Numbers and percentages indicate the number and proportion of proteins in each category. UP and DOWN refers to upregulated and downregulated proteins/genes, respectively. The data for Δ*suhB* were obtained by proteomics, while those for Δ*ecfG1*Δ*ecfG2* were obtained by transcriptomics.

We then examined whether any of the predicted SuhB targets listed in Table 1 corresponded to genes within the GSR regulon and showed the same expression pattern in Δ*suhB*, but none did (Figure 7). Although one predicted target (SGRAN_1462, an amidohydrolase) overlapped with the indirect GSR regulon, it was upregulated in the GSR deficient Δ*ecfG1*Δ*ecfG2* strain during stationary phase but downregulated in Δ*suhB* during exponential growth. Two additional targets, SGRAN_2291, encoding a probable DNA helicase II like protein, and SGRAN_2955, encoding a CheW protein, were downregulated in Δ*suhB* during stationary phase but upregulated in Δ*ecfG1*Δ*ecfG2* under the same conditions. In contrast, four predicted targets were upregulated in both mutants, though at different growth phases, exponential phase in Δ*suhB* and stationary phase in Δ*ecfG1*Δ*ecfG2*. These included ThnR (SGRAN_2804), a diacylglycerol O-acyltransferase (SGRAN_3397), a major facilitator superfamily protein (SGRAN_3990), and an alkyl sulfatase (SGRAN_4001). Overall, the SuhB regulon shows limited overlap with the GSR regulon. Only a small subset of genes exhibits concordant changes, being up or downregulated in both Δ*suhB* and the GSR regulon in the same growth phase, while most altered proteins are unique to each regulon. The concordant changes may only partially contribute to the stress phenotype observed in Δ*suhB*. Importantly, none of the predicted direct SuhB targets correspond to GSR regulatory genes nor to other genes within the GSR regulon, indicating that SuhB does not directly control the general stress response, and that the multiple stress-sensitivity phenotypes associated with *suhB* deletion do not result from alterations in the expression of GSR-regulated stress genes.

**Figure 7:**
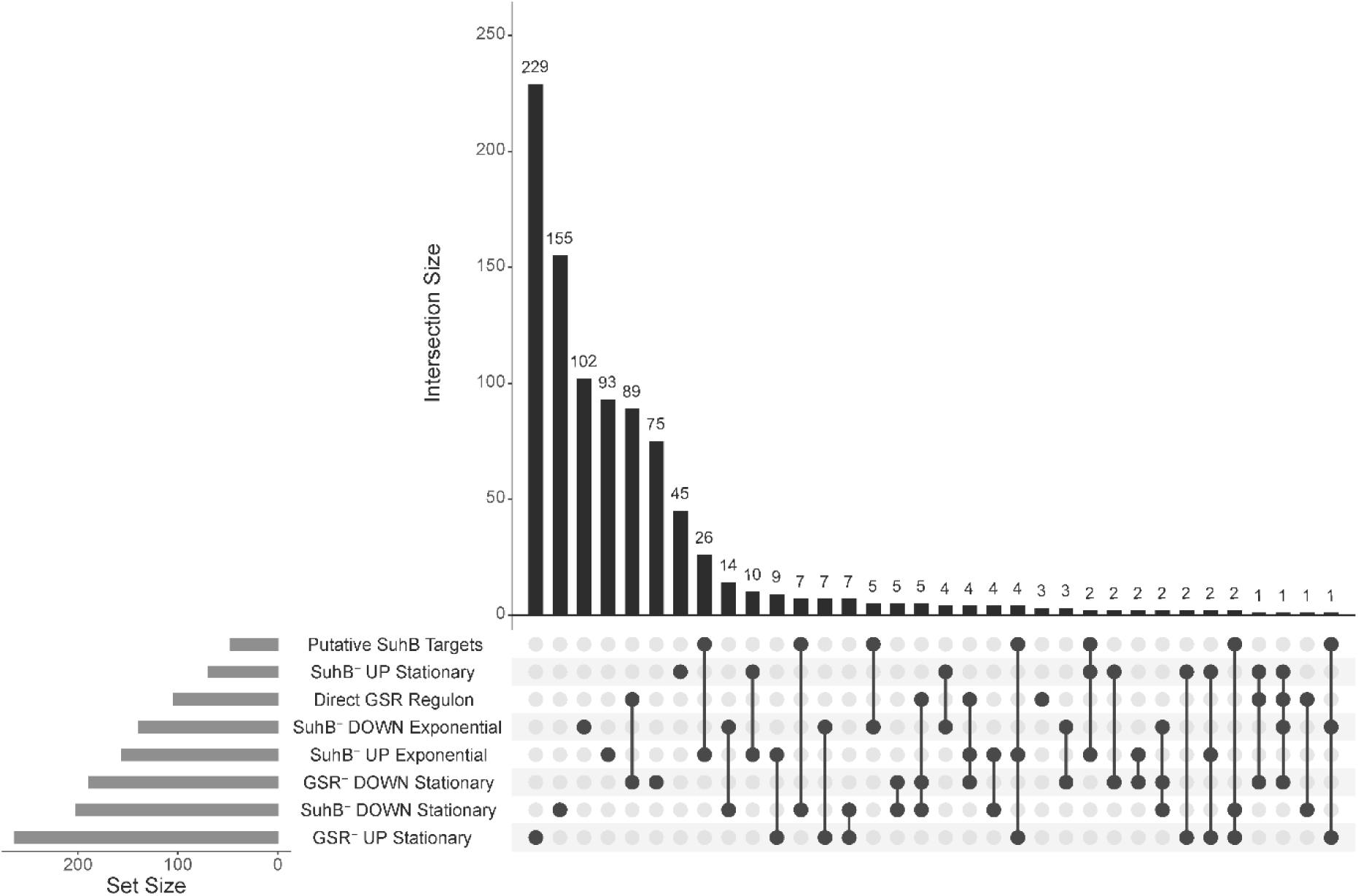
UpSet plot showing the overlap between SuhB regulated proteins and the GSR regulon. UpSet plot displaying the intersections among putative SuhB targets, proteins differentially expressed in the Δ*suhB* mutant under exponential or stationary growth (SuhB⁻ UP and SuhB⁻ DOWN), and genes belonging to the General Stress Response (GSR⁻ UP and GSR⁻ DOWN) regulon under stationary phase. The Direct GSR regulon refers to genes directly regulated by the EcfG sigma factors. Bars indicate the number of elements in each intersection (top) and in each individual dataset (left). The plot was constructed using the web server UpSetR (https://gehlenborglab.shinyapps.io/upsetr/) (*60*).

### SuhB expression is regulated by a LysR-type transcriptional regulator

To identify regulatory proteins that interact with the promoter region of the *suhB* gene, a promoter affinity chromatography assay was performed. A 308 bp DNA fragment containing the *suhB* promoter region was amplified by PCR and biotinylated at the 5′ end to allow its immobilization on streptavidin-coated magnetic beads. Protein extracts were prepared from *Sphingopyxis granuli* TFA cells grown in minimal medium (MM) supplemented either with sebacic acid, an inducing condition for *suhB* expression, or with tetralin, a non-inducing condition. Following the incubation of protein extracts with the immobilized DNA, the bound proteins were eluted and separated by SDS-PAGE. Bands that were differentially present between the inducing and non-inducing conditions were excised and analysed by MALDI-TOF mass spectrometry for protein identification (Supplementary Material. Figure S10). The proteins identified with the highest confidence scores included: a pyruvate carboxylase (band 1), propionyl-CoA carboxylase beta chain (SGRAN_2003; band 2), ATP-dependent RNA helicase RhlB (SGRAN_2295; band 3), pyruvate dehydrogenase E1 component subunit alpha (SGRAN_2408; band 4), a transcriptional regulator of the LysR family (SGRAN_2041; band 5), and the 30S ribosomal protein S3 (SGRAN_0303; band 6).

Among these, we focused our attention on the LysR-type transcriptional regulator (SGRAN_2041), as a potential direct regulator of *suhB* expression in TFA. LysR-type transcriptional regulators (LTTR) are part of a large protein family and display a well-conserved structure with an N-terminal HTH (helix-turn-helix) DNA-binding motif and a C-terminal effector domain. LTTR orthologues are known to contribute to all aspects of metabolism and physiology. This protein not only appeared as a specific interactor with the *suhB* promoter under inducing conditions, but also, sequence analysis revealed the presence of a putative LysR-binding site within the promoter region (Figure 8a).

**Figure 8:**
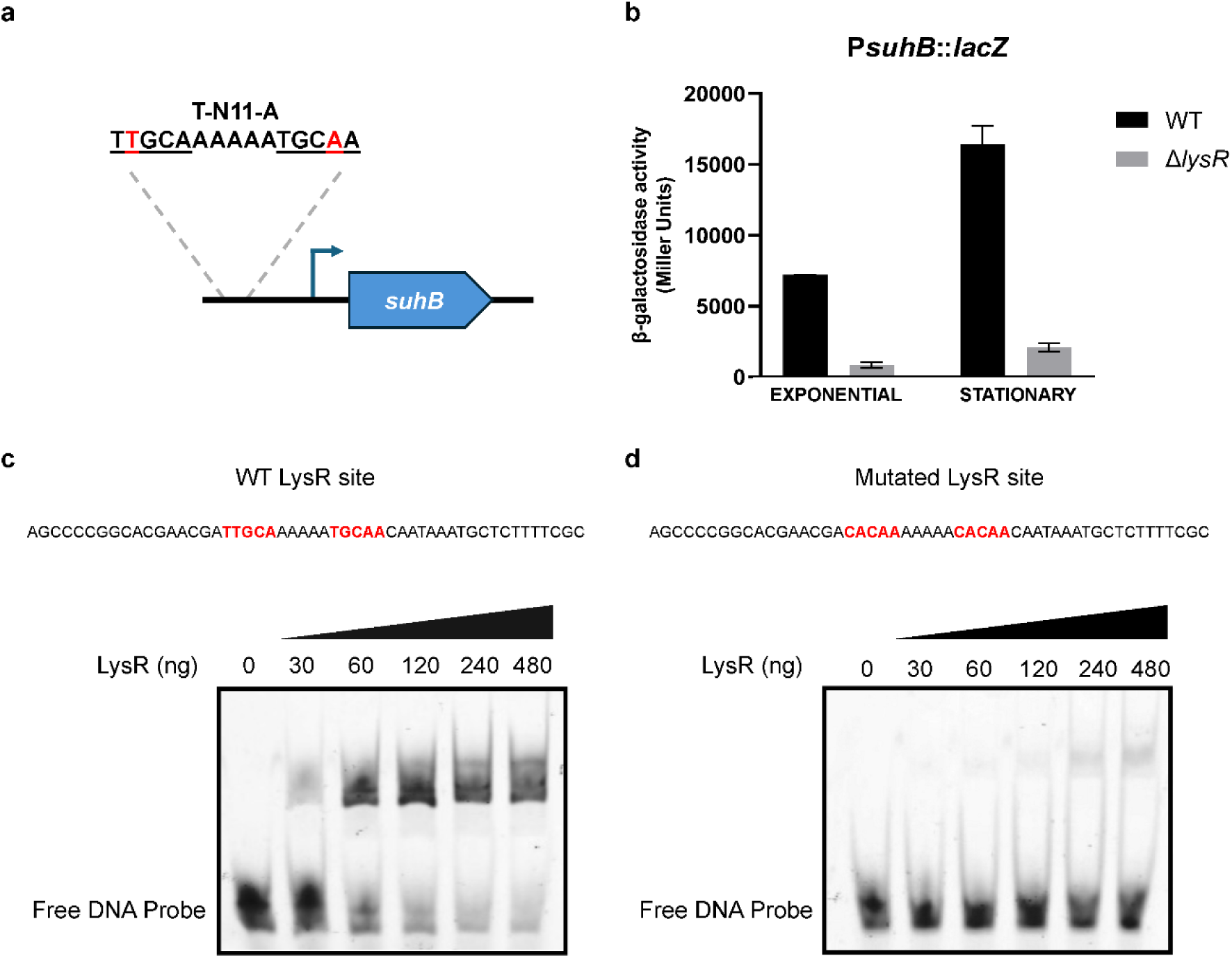
Analysis of the interaction between LysR and the *suhB* promoter region. (a) Schematic representation of the predicted LysR binding site (T–N₁₁–A motif) upstream of the *suhB* promoter. (b) β-galactosidase activity of the *PsuhB::lacZ* transcriptional fusion in WT and Δ*lysR* strains during exponential and stationary phases of growth in minimal medium (MM) supplemented with 40 mM BHB. (c) EMSA showing binding of LysR to a DNA probe containing the WT LysR binding site. Increasing amounts of LysR protein (0-480 ng) were added to the reactions. The shifted bands indicate formation of the LysR-DNA complex, whereas the free probe is observed at the bottom of the gel. (d) EMSA using a probe containing a mutated version of the LysR binding site.

Functional validation of the role of this regulator was carried out using an in-frame deletion mutant lacking SGRAN_2041 (strain MPO830). As expected, this mutant displayed increased sensitivity to all stress conditions tested compared to the wild-type strain (Figure 9). Moreover, the expression of *suhB*, quantified from a P*suhB::lacZ* fusion introduced into the chromosome of the LysR mutant, was significantly reduced in the mutant cultured in MM supplemented with 40 mM BHB, compared to the WT, further supporting the role of the LysR regulator as a transcriptional activator (Figure 8b).

**Figure 9:**
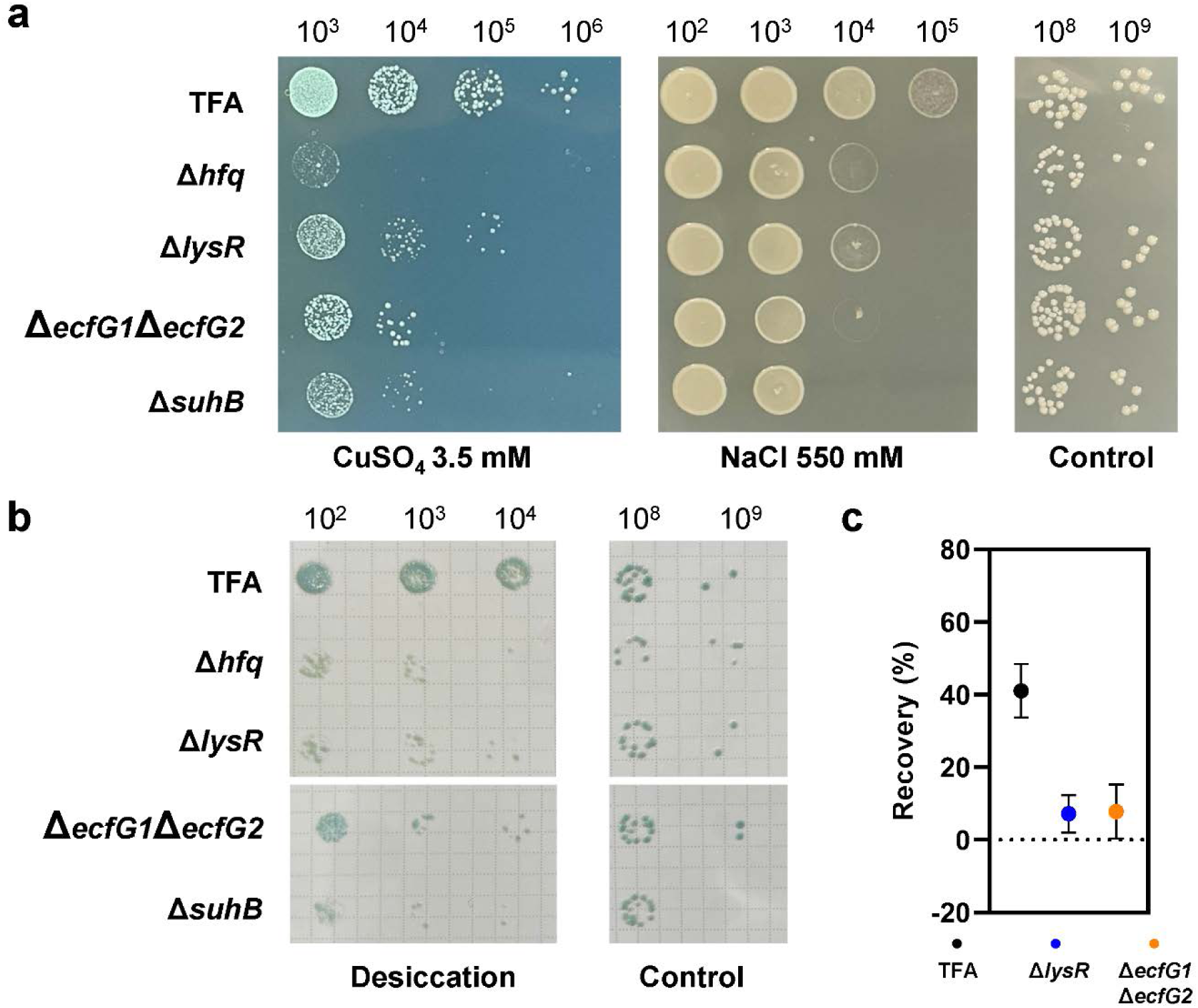
Phenotypic stress assays comparing *lysR* deletion mutant with the wild type TFA, mutant lacking the GSR sigma factors and *suhB* and *hfq* mutants. (a) Stress sensitivity was assessed by spotting serial dilutions of the different strains onto MML rich agar supplemented with 3.5 mM CuSO_4_ (heavy metal stress) or 500 mM NaCl (osmotic stress). (b) Desiccation tolerance was evaluated by placing serial dilutions on filters and allowing them to air dry for 24 h. (c) The ability to recover from oxidative stress was determined by comparing treated cultures to untreated controls 5 h after exposure to 10 mM H_2_O_2_.

To confirm direct interaction with the *suhB* promoter, electrophoretic mobility shift assays (EMSAs) were performed using purified LysR protein. A clear retarded band was observed when the promoter fragment was incubated with increasing concentrations of the regulator. This band shift was only detected when the predicted LysR-binding site was present in the DNA probe (Figure 8c). In contrast, when using a probe carrying a mutated LysR-binding site, the shift was almost undetectable (Figure 8d), further supporting the specificity of the interaction.

Together, these results strongly support that SGRAN_2041 encodes a LysR-type transcriptional regulator that directly binds the *suhB* promoter to activate its expression under inducing conditions.

## Discussion

Small RNAs (sRNAs) are key post-transcriptional regulators that modulate multiple biological functions in bacteria including stress response (*50*). One well characterized example is RyhB in *E. coli*. This sRNA is expressed under iron-limiting conditions, when the Fur repressor releases from its binding site in the *ryhB* promoter (*51, 52*). Additionally, RyhB has been shown to directly regulate more than 18 operons, and several additional genes have been identified as indirect targets. These findings highlight the major role of RyhB, together with the Fur protein, in the control of iron homeostasis (*53*). In this study, we demonstrated that SuhB, an small RNA that has been previously described to block the translation of the tetralin transcriptional activator ThnR (*31*), is also involved in multiple stress resistance in TFA. In absence of this sRNA, TFA is more sensitive to heavy metals, dissection, osmotic stress and oxidative stress in a similar way than the GSR impair mutant (*23*) (Figure 1). The proteomic changes observed between the WT and the SuhB deletion mutant, together with the predicted targets of this small RNA, support the notion that SuhB functions as a global regulator involved in the control of the stress response (among other possible functions yet unexplored). Several genes appear to be directly regulated by SuhB (Table 1), while others may represent indirect targets whose protein expression is also affected in the absence of this sRNA. Overall, in the SuhB deletion mutant, 156 proteins were upregulated and 139 downregulated during the exponential phase, whereas 69 proteins were upregulated and 202 downregulated during the stationary phase (where GSR is activated and stress condition is sensed), indicating a substantial alteration of the global proteome (Figure 2).

Among the functions upregulated in the Δ*suhB* mutant, we identified proteins involved in lipid metabolism as well as membrane-associated proteins, the latter showing a strong enrichment of TBDRs during the exponential phase. This pattern contrasts with what has been reported for *Caulobacter crescentus*, where overexpression of the sRNA CrfA induces TBDR expression (*54*). In *E. coli*, however, the Hfq-dependent sRNAs OmrA and OmrB repress the translation of various TBDRs, including CirA, FecA and FepA, as well as additional outer membrane proteins (*55*). Downregulation of such outer membrane channels and surface structures may be advantageous for the cell. For instance, it has been proposed that reducing TBDR abundance mitigates damage under osmotic stress by altering the interaction between the outer and inner membranes (*55*). In this context, the elevated abundance of multiple TBDRs observed in Δ*suhB* could contribute to the increased sensitivity of the mutant to osmotic stress. Notably, one of these receptors, SGRAN_4018, has been identified as a direct target of SuhB, further supporting the idea that this sRNA plays a role in tuning outer membrane permeability and maintaining envelope homeostasis. TonB systems require substantial energy to couple cytoplasmic membrane energy to outer membrane receptors (*56*). No differences in growth between the wild type and the Δ*suhB* mutant were observed under non-stress conditions (Figure 1, control). Nevertheless, the disruption of membrane homeostasis in the mutant is likely to compromise its ability to mount an effective stress response, which may partially explain the increased sensitivity of Δ*suhB* to multiple stressors. Although the SuhB regulon (stationary phase) is largely distinct from the GSR regulon (stationary phase) described in TFA (*23*), some stress related protein such as CspA (cold-shock protein) and SodB (superoxide dismutase) are downregulated in both the SuhB and the GSR deficient mutants (Δ*ecfG1*Δ*ecfG2*), suggesting that although SuhB and the GSR act through largely separate pathways, their regulons converge on certain genes that are critical for stress tolerance (*57, 58*). Neither CspA nor SodB has been predicted to be a direct target of SuhB, indicating that their decreased abundance is presumably an indirect regulatory effect.

Another important phenotype of Δ*suhB* is the overaccumulation of PHB in a similar way that Δ*mmgR* in *Sinorhizobium meliloti* (*30*). The Δ*suhB* strain appears to maximize its chances of survival under carbon excess conditions by redirecting acetyl-CoA toward the synthesis of this storage polymer. This metabolic shift is consistent with the observed upregulation of the acetoacetyl-CoA reductase PhbB (SGRAN_3755), identified in this study as a direct target of SuhB and overexpressed in its absence (Figure 4, Supplementary Data 1). In TFA, the Δ*suhB* mutant also accumulates a Phasin (SGRAN_1294) during exponential phase. Phasins are low molecular weight proteins associated with PHB granules, where they bind to the granule surface and promote PHB synthesis (*59*). Although this particular Phasin has not been predicted as a direct SuhB target, Phasins also accumulate in the Δ*mmgR* mutant of *Sinorhizobium meliloti*, where this effect has been proposed to occur either directly or indirectly regulated by MmgR (*30*).

We have also identified a LysR-type transcriptional regulator (LTTR), SGRAN_2041, that directly controls the transcription of *suhB*, thereby positioning this regulator upstream in the SuhB dependent stress response. The abundance of this regulator is not altered in the Δ*suhB* mutant (Supplementary Data 1) nor within the GSR regulon (*23*), suggesting that the observed effect on stress resistance may result from the reduced expression of SuhB in the absence of this LTTR. However, further studies on the regulon of this protein would be of interest to clarify whether it also controls additional stress-resistance mechanisms beyond the regulation of SuhB expression. Through its action as a small RNA, SuhB represses the translation of *thnR*, which encodes another LTTR that functions as the transcriptional activator of the tetralin degradation genes. This regulatory architecture establishes a hierarchical connection between both LysR regulators, whereby the LTTR upstream of SuhB modulates its expression, and SuhB then fine-tunes *thnR* translation.

In summary, our results highlight SuhB as a global regulator that contributes to stress resistance, envelope homeostasis, and metabolic regulation in TFA. The diverse changes observed in Δ*suhB* indicate that this sRNA influences multiple cellular processes, and further work will be needed to clarify the mechanisms behind these effects.

## Acknowledgements

This work was supported by grant PID2021-125491NB-I00 funded by MICIU/AEI/10.13039/501100011033 (Ministerio de Ciencia, Innovación y Universidades/Agencia Estatal de Investigación) and FEDER (EU) to F.R.-R. I.G.-R. was supported by a postdoctoral contract (PAIDI 2020, POSTDOC_21_00064) funded by the Andalusian Government (Junta de Andalucía). We are grateful to the members of the Division of Microbiology at the University Pablo de Olavide (Seville, Spain) for their advice and suggestions, and to the laboratory technicians for their invaluable technical support. We also thank the proteomics facility at CABD for their assistance. Finally, we thank Dr. Nadim Magdalani (Center for Cancer Research, National Cancer Institute, Bethesda, USA) for kindly providing the *E. coli* strain PM1805 and plasmid pNM46.

## References

1. G. Storz, J. Vogel, K. M. Wassarman, Regulation by small RNAs in bacteria: expanding frontiers. Mol Cell 43, 880–891 (2011).

2. A. Chauvier, N. G. Walter, Regulation of bacterial gene expression by non-coding RNA: It is all about time! Cell Chemical Biology 31, 71–85 (2024).

3. G. Storz, J. A. Opdyke, K. M. Wassarman, Regulating bacterial transcription with small RNAs. Cold Spring Harb Symp Quant Biol 71, 269–273 (2006).

4. J. Hör, G. Matera, J. Vogel, S. Gottesman, G. Storz, Trans-Acting Small RNAs and their effects on gene expression in *Escherichia coli* and *Salmonella enterica*. EcoSal Plus 9, (2020).

5. K. S. Fröhlich, K. Papenfort, Regulation outside the box: New mechanisms for small RNAs. Mol Microbiol 114, 363–366 (2020).

6. B. Felden, Y. Augagneur, Diversity and Versatility in Small RNA-Mediated Regulation in Bacterial Pathogens. Front Microbiol 12, 719977 (2021).

7. C. A. McCullen, J. N. Benhammou, N. Majdalani, S. Gottesman, Mechanism of positive regulation by DsrA and RprA small noncoding RNAs: pairing increases translation and protects rpoS mRNA from degradation. J Bacteriol 192, 5559–5571 (2010).

8. N. Majdalani, S. Chen, J. Murrow, K. St John, S. Gottesman, Regulation of RpoS by a novel small RNA: the characterization of RprA. Mol Microbiol 39, 1382–1394 (2001).

9. F. Repoila, S. Gottesman, Signal transduction cascade for regulation of RpoS: temperature regulation of DsrA. J Bacteriol 183, 4012–4023 (2001).

10. J. B. Rice, C. K. Vanderpool, The small RNA SgrS controls sugar-phosphate accumulation by regulating multiple PTS genes. Nucleic Acids Res 39, 3806–3819 (2011).

11. J. Borgmann, S. Schäkermann, J. E. Bandow, F. Narberhaus, A Small Regulatory RNA Controls Cell Wall Biosynthesis and Antibiotic Resistance. mBio 9, (2018).

12. Y. Han et al., Small RNA-regulated expression of eflux pump affects tigecycline resistance and heteroresistance in clinical isolates of Klebsiella pneumoniae. Microbiol Res 287, 127825 (2024).

13. C. Baussier, C. Oriol, S. Durand, B. Py, P. Mandin, Small RNA OxyS induces resistance to aminoglycosides during oxidative stress by controlling Fe-S cluster biogenesis in. Proc Natl Acad Sci U S A 121, e2317858121 (2024).

14. M. J. Hernáez, W. Reineke, E. Santero, Genetic analysis of biodegradation of tetralin by a Sphingomonas strain. Appl Environ Microbiol 65, 1806–1810 (1999).

15. I. García-Romero, J. Nogales, E. Díaz, E. Santero, B. Floriano, Understanding the metabolism of the tetralin degrader Sphingopyxis granuli strain TFA through genome-scale metabolic modelling. Sci Rep 10, 8651 (2020).

16. B. Floriano, E. Santero, F. Reyes-Ramírez, Biodegradation of Tetralin: Genomics, Gene Function and Regulation. Genes (Basel) 10, (2019).

17. I. García-Romero et al., Genomic analysis of the nitrate-respiring Sphingopyxis granuli (formerly Sphingomonas macrogoltabida) strain TFA. BMC Genomics 17, 93 (2016).

18. A. Francez-Charlot, A. Kaczmarczyk, H. M. Fischer, J. A. Vorholt, The general stress response in Alphaproteobacteria. Trends Microbiol 23, 164–171 (2015).

19. R. Hengge. (2011).

20. S. Bouillet, T. S. Bauer, S. Gottesman, RpoS and the bacterial general stress response. Microbiol Mol Biol Rev 88, e0015122 (2024).

21. R. de Dios, E. Santero, F. Reyes-Ramírez, Extracytoplasmic Function σ Factors as Tools for Coordinating Stress Responses. Int J Mol Sci 22, (2021).

22. R. de Dios, E. Santero, F. Reyes-Ramírez, The functional differences between paralogous regulators define the control of the general stress response in Sphingopyxis granuli TFA. Environ Microbiol 24, 1918–1931 (2022).

23. R. de Dios, E. Rivas-Marin, E. Santero, F. Reyes-Ramírez, Two paralogous EcfG σ factors hierarchically orchestrate the activation of the General Stress Response in Sphingopyxis granuli TFA. Sci Rep 10, 5177 (2020).

24. B. A. Berghoff, J. Glaeser, C. M. Sharma, J. Vogel, G. Klug, Photooxidative stress-induced and abundant small RNAs in Rhodobacter sphaeroides. Mol Microbiol 74, 1497–1512 (2009).

25. C. del Val et al., A survey of sRNA families in α-proteobacteria. RNA Biol 9, 119–129 (2012).

26. R. Madhugiri et al., Small RNAs of the Bradyrhizobium/Rhodopseudomonas lineage and their analysis. RNA Biol 9, 47–58 (2012).

27. N. G. Assis et al., Identification of Hfq-binding RNAs in Caulobacter crescentus. RNA Biol 16, 719–726 (2019).

28. K. S. Fröhlich, K. U. Förstner, Z. Gitai, Post-transcriptional gene regulation by an Hfq-independent small RNA in Caulobacter crescentus. Nucleic Acids Res 46, 10969–10982 (2018).

29. L. N. Vogt et al., Genome-wide profiling of Hfq-bound RNAs reveals the iron-responsive small RNA RusT in *Caulobacter crescentus*. mBio 15, e0315323 (2024).

30. A. Lagares, G. C. Borella, U. Linne, A. Becker, C. Valverde, Regulation of Polyhydroxybutyrate Accumulation in Sinorhizobium meliloti by the T*rans-*Encoded Small RNA MmgR. J Bacteriol 199, (2017).

31. I. García-Romero, K. U. Förstner, E. Santero, B. Floriano, SuhB, a small non-coding RNA involved in catabolite repression of tetralin degradation genes in Sphingopyxis granuli strain TFA. Environ Microbiol 20, 3671–3683 (2018).

32. X. Luo, N. Majdalani, Directed Screening for sRNA Targets in E. coli Using a Plasmid Library. Methods Mol Biol 2741, 291–306 (2024).

33. I. García-Romero, R. de Dios, F. Reyes-Ramírez, An improved genome editing system for Sphingomonadaceae. Access Microbiol 6, (2024).

34. Y. Hsiao et al., Analysis and Visualization of Quantitative Proteomics Data Using FragPipe-Analyst. J Proteome Res, (2024).

35. L. Tomás-Gallardo, E. Santero, B. Floriano, Involvement of a putative cyclic amp receptor protein (CRP)-like binding sequence and a CRP-like protein in glucose-mediated catabolite repression of thn genes in Rhodococcus sp. strain TFB. Appl Environ Microbiol 78, 5460–5462 (2012).

36. Y. W. Hsieh, A. Alqadah, C. F. Chuang, An Optimized Protocol for Electrophoretic Mobility Shift Assay Using Infrared Fluorescent Dye-labeled Oligonucleotides. J Vis Exp, (2016).

37. J. H. Miller, Experiments in Molecular Genetics. C. S. H. Laboratory., Ed., (New York, USA, 1972).

38. P. V. Sola et al., PHB in cyanobacteria: analyzing production through images processing and FT-IR techniques. N Biotechnol 89, 119–129 (2025).

39. S. H. Yu, J. Vogel, K. U. Förstner, ANNOgesic: a Swiss army knife for the RNA-seq based annotation of bacterial/archaeal genomes. Gigascience 7, (2018).

40. P. R. Wright et al., CopraRNA and IntaRNA: predicting small RNA targets, networks and interaction domains. Nucleic Acids Res 42, W119–123 (2014).

41. H. Tafer, I. L. Hofacker, RNAplex: a fast tool for RNA-RNA interaction search. Bioinformatics 24, 2657–2663 (2008).

42. U. Mückstein et al., Thermodynamics of RNA-RNA binding. Bioinformatics 22, 1177–1182 (2006).

43. R. Lorenz et al., ViennaRNA Package 2.0. Algorithms Mol Biol 6, 26 (2011).

44. A. Alexa, J. Rahnenführer, T. Lengauer, Improved scoring of functional groups from gene expression data by decorrelating GO graph structure. Bioinformatics 22, 1600–1607 (2006).

45. H. Wickham, ggplot2: Elegant Graphics for Data Analysis. Springer-Verlag New York, (2016).

46. A. Arce-Rodríguez, B. Calles, P. I. Nikel, V. de Lorenzo, The RNA chaperone Hfq enables the environmental stress tolerance super-phenotype of Pseudomonas putida. Environ Microbiol 18, 3309–3326 (2016).

47. K. Tang, N. Jiao, K. Liu, Y. Zhang, S. Li, Distribution and functions of TonB-dependent transporters in marine bacteria and environments: implications for dissolved organic matter utilization. PLoS One 7, e41204 (2012).

48. G. Martín-Cabello, E. Moreno-Ruiz, V. Morales, B. Floriano, E. Santero, Involvement of poly(3-hydroxybutyrate) synthesis in catabolite repression of tetralin biodegradation genes in Sphingomonas macrogolitabida strain TFA. Environ Microbiol Rep 3, 627–631 (2011).

49. J. C. Oliveros. (2007-2015).

50. S. Gottesman et al., Small RNA regulators and the bacterial response to stress. Cold Spring Harb Symp Quant Biol 71, 1–11 (2006).

51. E. Massé, S. Gottesman, A small RNA regulates the expression of genes involved in iron metabolism in Escherichia coli. Proc Natl Acad Sci U S A 99, 4620–4625 (2002).

52. E. Holmqvist, E. G. H. Wagner, Impact of bacterial sRNAs in stress responses. Biochem Soc Trans 45, 1203–1212 (2017).

53. E. Massé, C. K. Vanderpool, S. Gottesman, Effect of RyhB small RNA on global iron use in Escherichia coli. J Bacteriol 187, 6962–6971 (2005).

54. S. G. Landt, J. A. Lesley, L. Britos, L. Shapiro, CrfA, a small noncoding RNA regulator of adaptation to carbon starvation in Caulobacter crescentus. J Bacteriol 192, 4763–4775 (2010).

55. M. Guillier, S. Gottesman, Remodelling of the Escherichia coli outer membrane by two small regulatory RNAs. Mol Microbiol 59, 231–247 (2006).

56. Q. Zhao et al., Influence of the TonB energy-coupling protein on eflux-mediated multidrug resistance in Pseudomonas aeruginosa. Antimicrob Agents Chemother 42, 2225–2231 (1998).

57. C. S. Bakshi et al., Superoxide dismutase B gene (sodB)-deficient mutants of Francisella tularensis demonstrate hypersensitivity to oxidative stress and attenuated virulence. J Bacteriol 188, 6443–6448 (2006).

58. B. Schmid et al., Role of cold shock proteins in growth of Listeria monocytogenes under cold and osmotic stress conditions. Appl Environ Microbiol 75, 1621–1627 (2009).

59. C. Wang et al., Influence of the poly-3-hydroxybutyrate (PHB) granule-associated proteins (PhaP1 and PhaP2) on PHB accumulation and symbiotic nitrogen fixation in Sinorhizobium meliloti Rm1021. J Bacteriol 189, 9050–9056 (2007).

60. A. Lex, N. Gehlenborg, H. Strobelt, R. Vuillemot, H. Pfister, UpSet: Visualization of Intersecting Sets. IEEE Trans Vis Comput Graph 20, 1983–1992 (2014).

