## Supplementary Materials for "Uncovering the regulatory network of the small RNA SuhB and its contribution to stress resistance in *Sphingopyxis granuli* TFA"

[1]Centro Andaluz de Biología del Desarrollo, Universidad Pablo de Olavide/Consejo Superior de Investigaciones Científicas/Junta de Andalucía, ES-41013 Seville, Spain

[2]Departamento de Biología Molecular e Ingeniería Bioquímica, Universidad Pablo de Olavide, ES-41013 Seville, Spain.

**Supplementary Table 1**: Antibiotic concentration for *E. coli* and *S. granuli*

| Specie | Antibiotic | Concentration (µg/mL) |
|---|---|---|
| ***E. coli*** | Kanamycin | 25 |
| | Ampicilin | 100 |
| | Clroramphenicol | 15 |
| ***S. granuli*** | Kanamycin | 20 |
| | Ampicilin | 5 |
| | Streptomycin | 50 or 200 |
| | Tetracycline | 10 |

**Supplementary Table 2**: Strains and plasmids used in this study.

| Strains | Description | Reference |
|---|---|---|
| ***Escherichia coli*** | | |
| **DH5α** | F- ø80lacZΔM15 ΔlacZYA-argF U169 recA1 endA1 hsdR17(rK-mK-) | (*1*) |
| **DH5α λ*pir*** | λ*pir* phage lysogen of DH5α | Lab collection |
| **PM1805** | MG1655 *mal::lacI*$^q$ *ΔaraBAD araC*$^+$ *lacI'::*P$_{BAD}$-*cat-sacB-lacZ*, mini-λtetR *trpE*$^+$ | (*2*) |
| **MPO667** | Derived from PM1805. MG1655 *mal::lacI*$^q$ *ΔaraBAD araC*$^+$ *lacI'::*P$_{BAD}$-*cat-SGRAN_4018-lacZ trpE*$^+$ | This work |
| **MPO668** | Derived from PM1805. MG1655 *mal::lacI*$^q$ *ΔaraBAD araC*$^+$ *lacI'::*P$_{BAD}$-*cat-SGRAN_3755-lacZ trpE*$^+$ | This work |
| ***Sphingopyxis granuli*** | | |
| **TFA** | Wild type. Str$^r$. | (*3*) |
| **MPO209** | MiniTn5Km Insertion in PHA Synthase. Str$^r$ Km$^r$ | (*4*) |
| **MPO641** | *PsuhB::lacZ* transcriptional fusion integrated into TFA chromosome. Str$^r$ Ap$^r$. | (*5*) |
| **MPO642** | Substitution of *suhB* gene by a kanamycin resistance gene. Str$^r$ Km$^r$. | (*5*) |
| **MPO646** | *hfq* deletion mutant. Str$^r$. | This work |
| **MPO830** | SGRAN_2041 (*lysR*) deletion mutant. Str$^r$. | This work |
| **MPO831** | *PsuhB::lacZ* transcriptional fusion integrated into MPO830 chromosome. Str$^r$ Ap$^r$. | This work |
| **MPO858** | *nepR2::lacZ* translational fusion integrated into TFA chromosome. Str$^r$ Ap$^r$. | (*6*) |
| **MPO860** | *ecfG1* and *ecfG2* deletion mutant. Str$^r$. | (*6*) |
| **MPO939** | *nepR2::lacZ* translational fusion integrated into MPO642 chromosome. Str$^r$ Ap$^r$ Km$^r$. | This work |
| **Plasmids** | **Description** | **Reference** |
| **pLAFR3** | Broad spectrum cosmid vector. Mob$^+$, Tra$^-$, Tc$^r$ | (*7*) |
| **pMPO1408** | pJES379 derivative bearing a *nepR2::lacZ* translational fusion, Ap$^r$. | (*6*) |
| **pMPO1412** | pEMG derivative with *rpsL1* streptomycin counterselection marker, Km$^r$. | (*6, 8*) |
| **pMPO1152** | *suhB* gene cloned into pLAFR3 under its own promoter , Tc$^r$ | (*5*) |
| **pMPO1151** | *PsuhB::lacZ* transcriptional fusion cloned into pIC552, Ap$^r$. | (*5*) |
| **pMPO1158** | pSEVA224 derivative bearing *hfq* expressed under P$_{trc}$ promoter, Km$^r$. | This work |
| **pMPO1159** | pSEVA221 derivative bearing *hfq* expressed under its own promoter, Km$^r$ | This work |
| **pMPO1161** | pMPO1412 derivative bearing *hfq* flanking regions for its deletion, Km$^r$. | This work |

| | | |
|---|---|---|
| **pMPO1169** | pTYB21 derivative bearing *intein*::SGRAN_2041 (*lysR*) fusion for LysR purification, Ap$^r$. | This work |
| **pMPO1178** | pNM46 derivative bearing *suhB* expressed under $P_{lac}$ promoter, Ap$^r$. | This work |
| **pMPO1822** | pMPO1412 derivative bearing SGRAN_2041 (*lysR*) flanking regions for its deletion, Km$^r$. | This work |
| **pNM46** | pBR-plac carrying the *lacI* gene, Ap$^r$. | (*2*) |
| **pSEVA221** | oriRK2, standard polylinker, Km$^r$ | (*9*) |
| **pSEVA224** | oriRK2, standard polylinker, *lacI*$^q$-$P_{trc}$, Km$^r$. | (*9*) |
| **pSW-I** | oriRK2, xylS, $P_m$::*I-sceI*, Ap$^r$. | (*10*) |
| **pTYB21** | *E. coli* expression vector containing an N-terminal fusion intein variant for protein purification using the IMPACT kit, Ap$^r$. | New England Biolabs |

**Supplementary Table 3**: Oligonucleotides used in this study.

| **Oligonucleotide** | **Sequence** | **Use** |
|---|---|---|
| **BamHI_hfq_rv** | ATATAGGATCCTCAGTCGCCGGAGTCGCCTTC | PCR of *hfq* |
| **EcoRI_tsshfq_fw** | ATATAGAATTCGCGAAGATGCAGGCTGCCGG | PCR of *hfq* |
| **F24** | CGCCAGGGTTTTCCCAGTCACGAC | PCR and sequencing of pSEVA224 insert |
| **pSEVA224_seq_F** | TCATCCGGCTCGTATAATGTGTGG/ | PCR and sequencing of pSEVA224 insert |
| **hfq_up_EcoRI_F** | ATATAGAATTCTCATTTCGTGCGGCGTTATGC | PCR of upstream region of *hfq* |
| **hfq_up_HindIII_R** | ATATAAAGCTTATCGGACACATTGGCCTCCTG | PCR of upstream region of *hfq* |
| **hfq_down_HindIII_F** | ATATAAAGCTTGGCGACTGATCCGATGGACGA | PCR of downstream region of *hfq* |
| **hfq_down_XbaI_R** | ATATATCTAGAGGTGTCGATCTTGTTCCATATC | PCR of downstream region of *hfq* |
| **check_hfq_mut_FW** | GGAACATCCCGCCTTTTCC | PCR and sequencing to checking *hfq* deletion |
| **check_hfq_mut_RV** | GATGCCGGGCTTGATCGC | PCR and sequencing to checking *hfq* deletion |
| **pBR-Rv** | GCATTGTTAGATTTCATACACGG | To verify and sequence insert in pNM46 |
| **pBR-Fw** | GGAAAACGTTCTTCGGGGCG | To verify and sequence insert in pNM46 |
| **AatII_suhB_Fw** | ATATAGACGTCCACACTTGCCTTTTAGGCATCCTCTC | PCR of *suhB* |
| **EcoRI_suhB_Rv** | ATATAGAATTCGGGCAATTCGGCACGGACGAAAAAAAAGG | PCR of *suhB* |
| **pBAD.for** | CCACATTGATTATTTGCACGGCG | To amplify and sequence allelic |

| | | |
|---|---|---|
| | | exchange in the PM1805 strain |
| **PM1805_lacZ_Rv** | GCAACTGTTGGGAAGGGCGATC | To amplify and sequence allelic exchange in the PM1805 strain |
| **SGRAN_2041_ATG_F** | ATGCAACGCCTTCCCCCCCTC | PCR of SGRAN_2041 |
| **SGRAN_2041_BamHI_R** | ATATAGGATCCTCAGATTTTCGTCGCCAGC | PCR of SGRAN_2041 |
| **PM1805_3755_F** | ACCTGACGCTTTTTATCGCAACTCTCTACTG TTTCTCCATAGAACATCTGTGGGAGAGCAAC | For allelic replacement of the cat-sacB genes in PM1805 with the first 25 aa of SGRAN_3755 in phase with LacZ |
| **PM1805_3755_R** | TAACGCCAGGGTTTTCCCAGTCACGACGTT GTAAAACGACCATGCCCTGCAGCTTCAGCG | For allelic replacement of the cat-sacB genes in PM1805 with the first 25 aa of SGRAN_3755 in phase with LacZ |
| **PM1805_4018_F** | ACCTGACGCTTTTTATCGCAACTCTCTACTG TTTCTCCATACTCATTCGAGAATGTGACG | For allelic replacement of the cat-sacB genes in PM1805 with the first 25 aa of SGRAN_4018 in phase with LacZ |
| **PM1805_4018_R** | TAACGCCAGGGTTTTCCCAGTCACGACGTT GTAAAACGACGACCTGCGCCATGGCCGAGA C | For allelic replacement of the cat-sacB genes in PM1805 with the first 25 aa of SGRAN_4018 in phase with LacZ |
| **Long_LysR_suhB_F** | AGCCCCGGCACGAACGATTGCAAAAAATGC AACAATAAATGCTCTTTTCGC | To construct the WT probe of the SuhB promoter |
| **LysR_suhB_mut_F** | AGCCCCGGCACGAACGACACAAAAAAACAC AACAATAAATGCTCTTTTCGC | To construct the mutated probe of the SuhB promoter |
| **Dye_lysR_suhB** | [6FAM]GCGAAAAGAGCATT | To construct the WT and mutated probe of the SuhB promoter. Labelled with 6FAM |
| **LysRSuhBDwn_BF_F** | AAAAGGATCCCGTTGCATCATCGTCTCCT | PCR of downstream region of SGRAN_2041 |
| **LysRSuhBUp_BF_F** | AAAAGAATTCATCCGACCGAAGCGAAAC | PCR of upstream region of SGRAN_2041 |
| **LysRSuhBDwn_BF_R** | AAAATCTAGAGGCCGTTGATCAGGAAGAAT | PCR of downstream region of SGRAN_2041 |

| | | |
|---|---|---|
| **LysRSuhBUp_BF_R** | AAAAGGATCCGTGAAGCTGTTCCACGACTG | PCR of downstream region of SGRAN_2041 |
| **suhB_for** | TCGTCCCAATCTAGGCGATA | PCR of SuhB promoter |
| **suhB_rev_Btn** | ATATATGCCGCCGGATTTTC | PCR of SuhB promoter. Biotinylated |
| **KmFw-pk18** | GATTGAACAAGATGGATTGC | PCR to check first integration of pMPO1412 derivative plasmids |
| **KmRev-pK18** | CGTCAAGAAGGCGATAGAAGG | PCR to check first integration of pMPO1412 derivative plasmids |

**Supplementary Table 4**: Abundance of GSR regulatory proteins in Δ*suhB* compared to WT in exponential and stationary phase. Log2(FC) should be > 1 or < -1 and adjusted p-value lower than 0.05 to be significant.

| Protein ID | Log2(FC) Exponential / Adjusted p-value | Log2(FC) Stationary / Adjusted p-value | Significant |
|---|---|---|---|
| EcfG1 (SGRAN_1161) | ND / ND | -0.326 / 0.742 | No |
| EcfG2 (SGRAN_1163) | 0.378 / 0.367 | -2.54 / 0.0868 | No |
| NepR1 (SGRAN_0992) | -0.975 / 0.193 | -0.299 / 0.756 | No |
| NepR2 (SGRAN_1162) | ND / ND | ND / ND | -- |
| PhyR1 (SGRAN_0993) | -0.274 / 0.605 | -3.56 / 0.088 | No |
| PhyR2 (SGRAN_1164) | -0.116 / 0.868 | -1.56 / 0.365 | No |

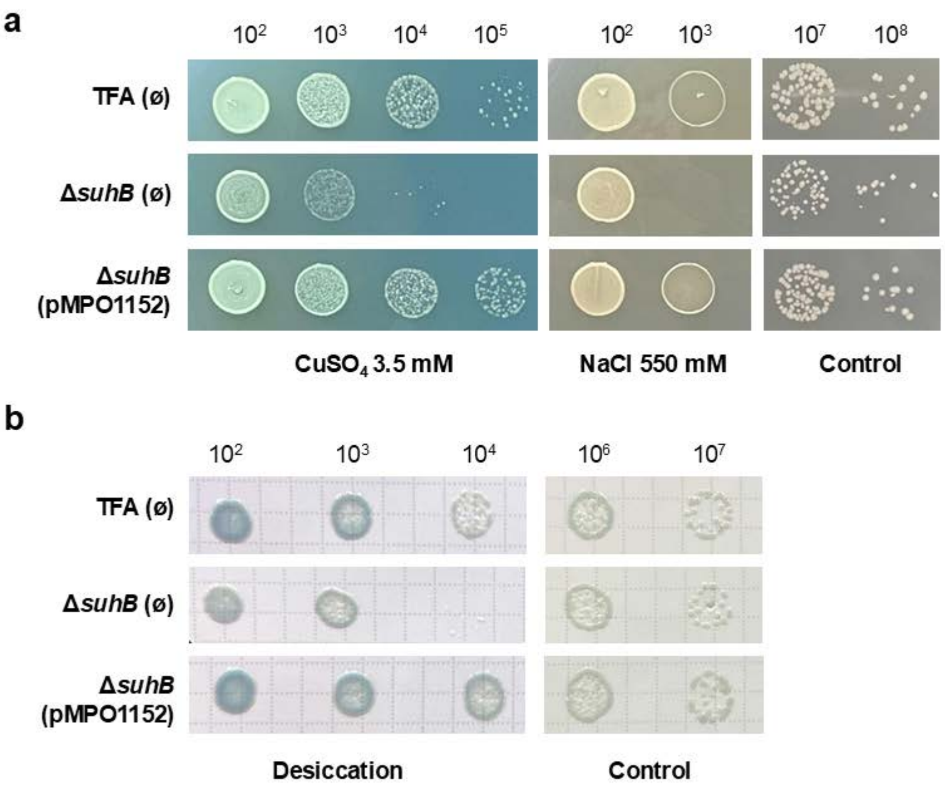


**Supplementary Figure 1: Phenotypic stress assays comparing the WT strain, the *suhB* deletion mutant carrying the empty plasmid (pLAFR3), and the mutant complemented with *suhB* expressed from its own promoter (pMPO1152).** (a) Stress sensitivity was assessed by spotting serial dilutions of the different strains onto MML rich agar supplemented with 3.5 mM $CuSO_4$ (heavy metal stress) or 500 mM NaCl (osmotic stress). (b) Desiccation tolerance was evaluated by placing serial dilutions on filters and allowing them to air dry for 24 h.

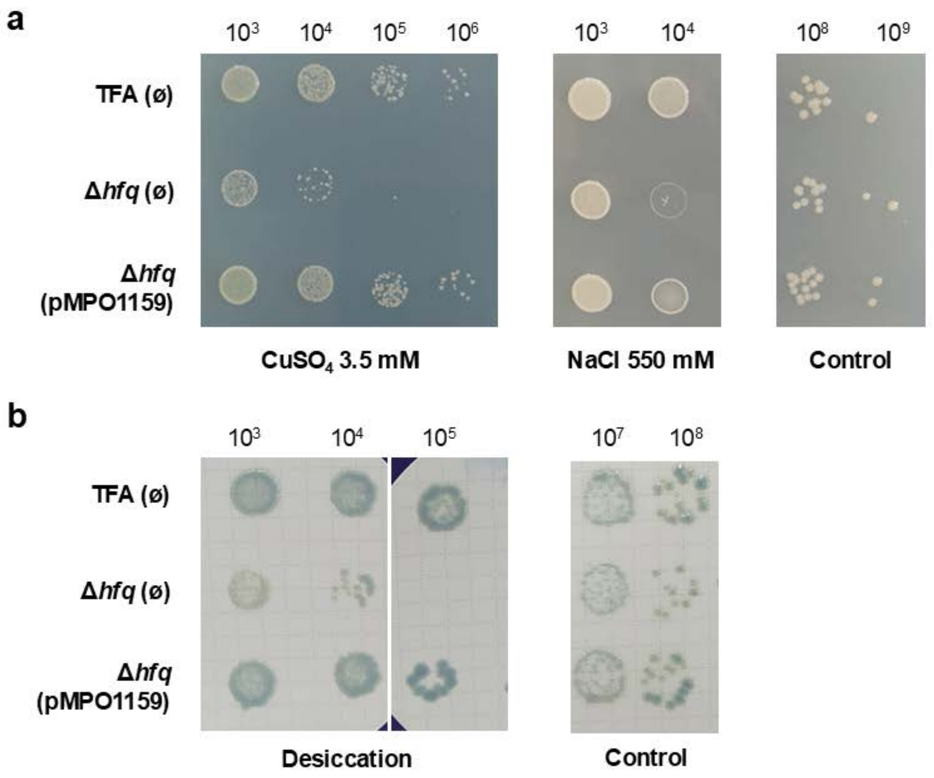


**Supplementary Figure 2: Phenotypic stress assays comparing the WT strain, the *hfq* deletion mutant carrying the empty plasmid (pSEVA221), and the mutant complemented with *hfq* expressed from its own promoter (pMPO1159).** (a) Stress sensitivity was assessed by spotting serial dilutions of the different strains onto MML rich agar supplemented with 3.5 mM $CuSO_4$ (heavy metal stress) or 500 mM NaCl (osmotic stress). (b) Desiccation tolerance was evaluated by placing serial dilutions on filters and allowing them to air dry for 24 h.

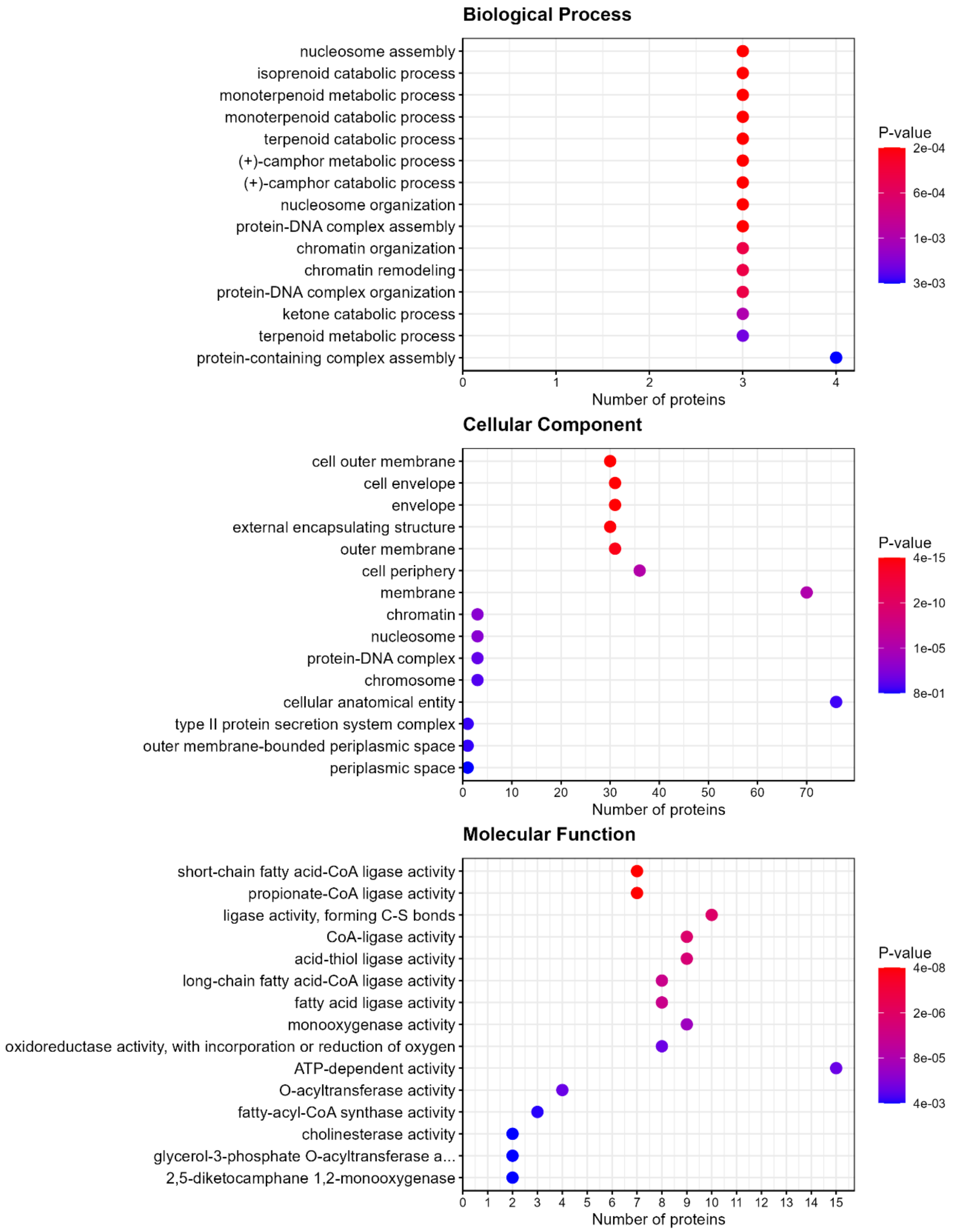


**Supplementary Figure 3: GO term enrichment of proteins upregulated in Δ*suhB* relative to the WT during exponential phase.** The X-axis shows the number of genes per category, and bar color indicates the p-value, with redder bars representing lower p-values.

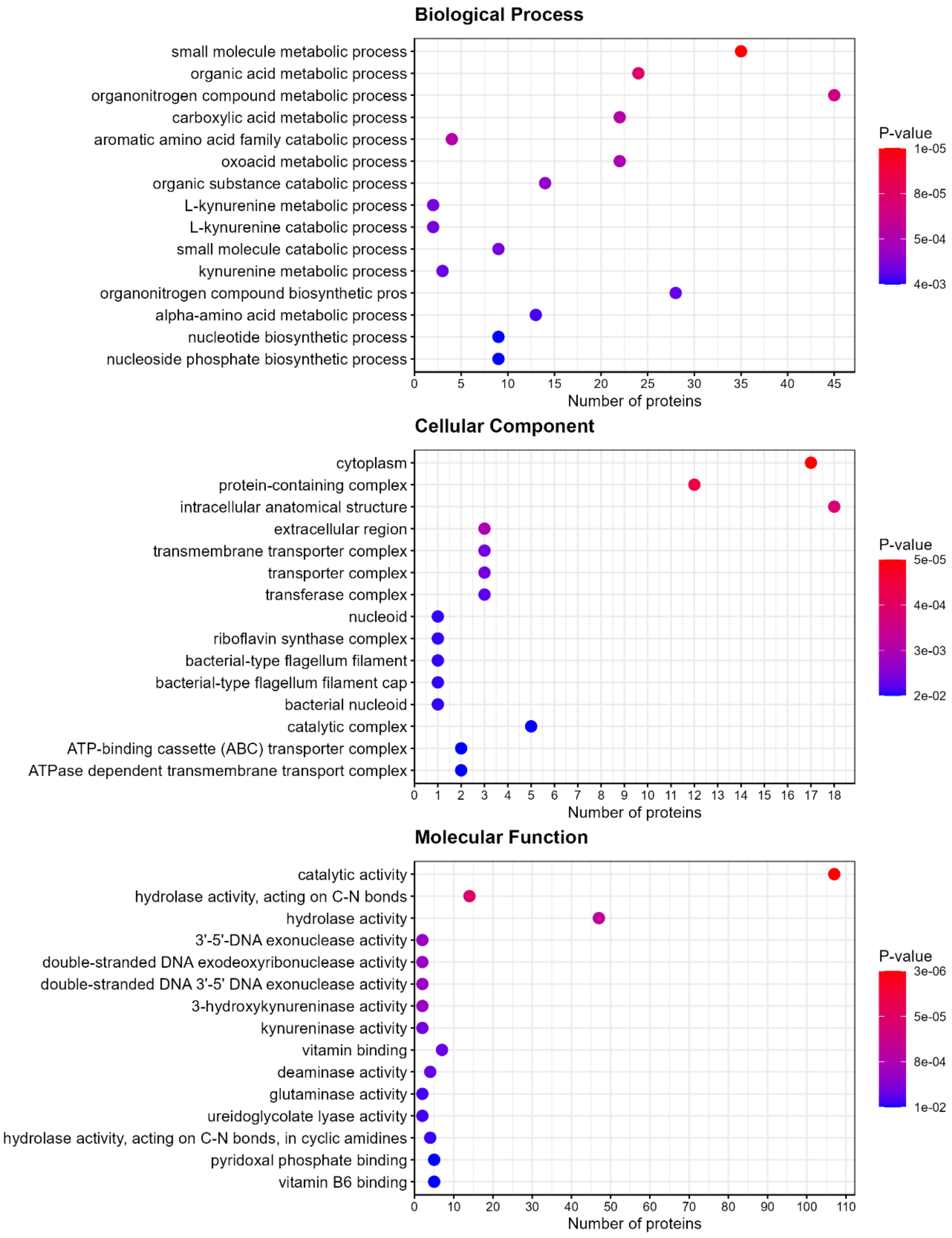


**Supplementary Figure 4: GO term enrichment of proteins downregulated in Δ*suhB* relative to the WT during exponential phase.** The X-axis shows the number of genes per category, and bar color indicates the p-value, with redder bars representing lower p-values.

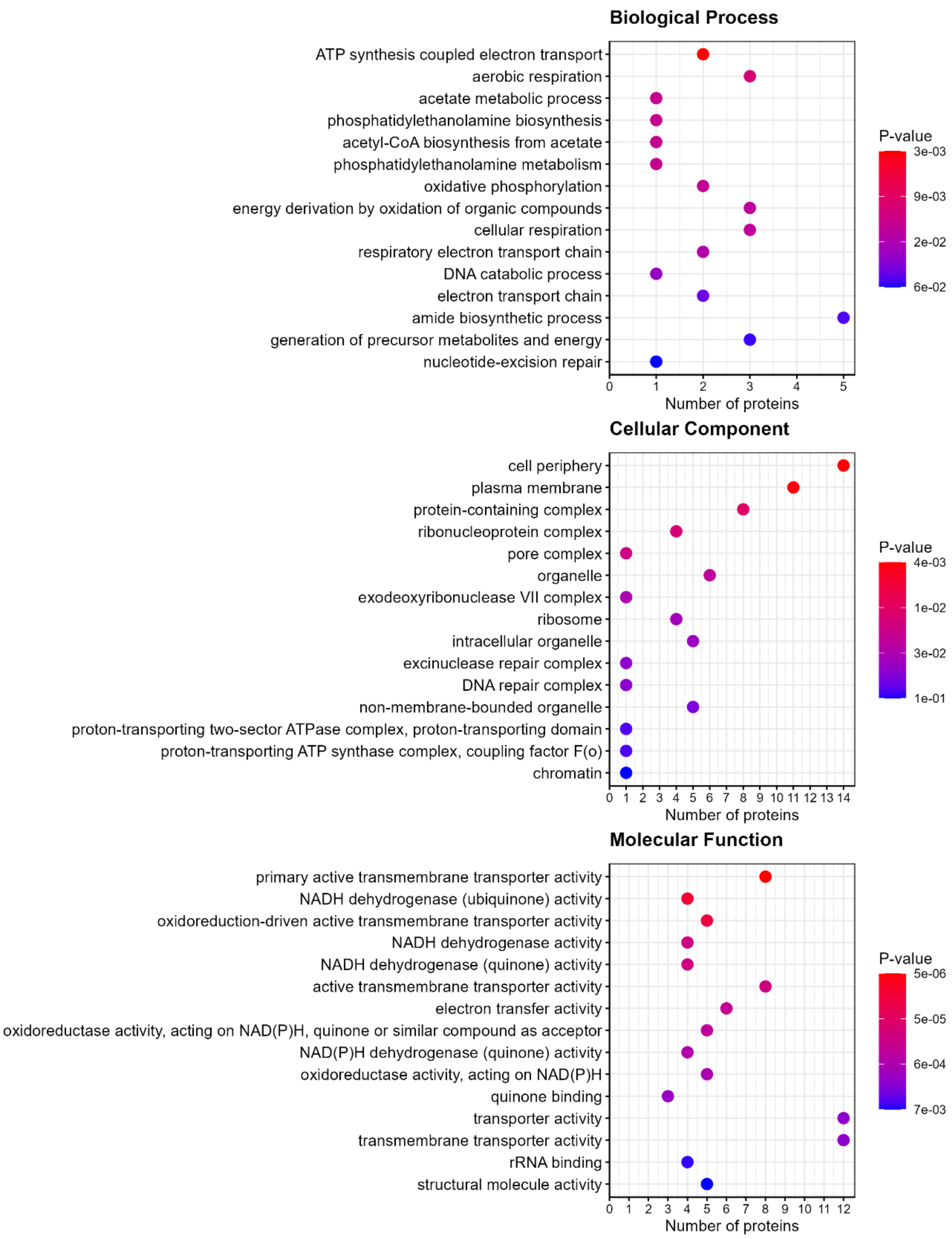


**Supplementary Figure 5: GO term enrichment of proteins upregulated in Δ*suhB* relative to the WT during stationary phase.** The X-axis shows the number of genes per category, and bar color indicates the p-value, with redder bars representing lower p-values.

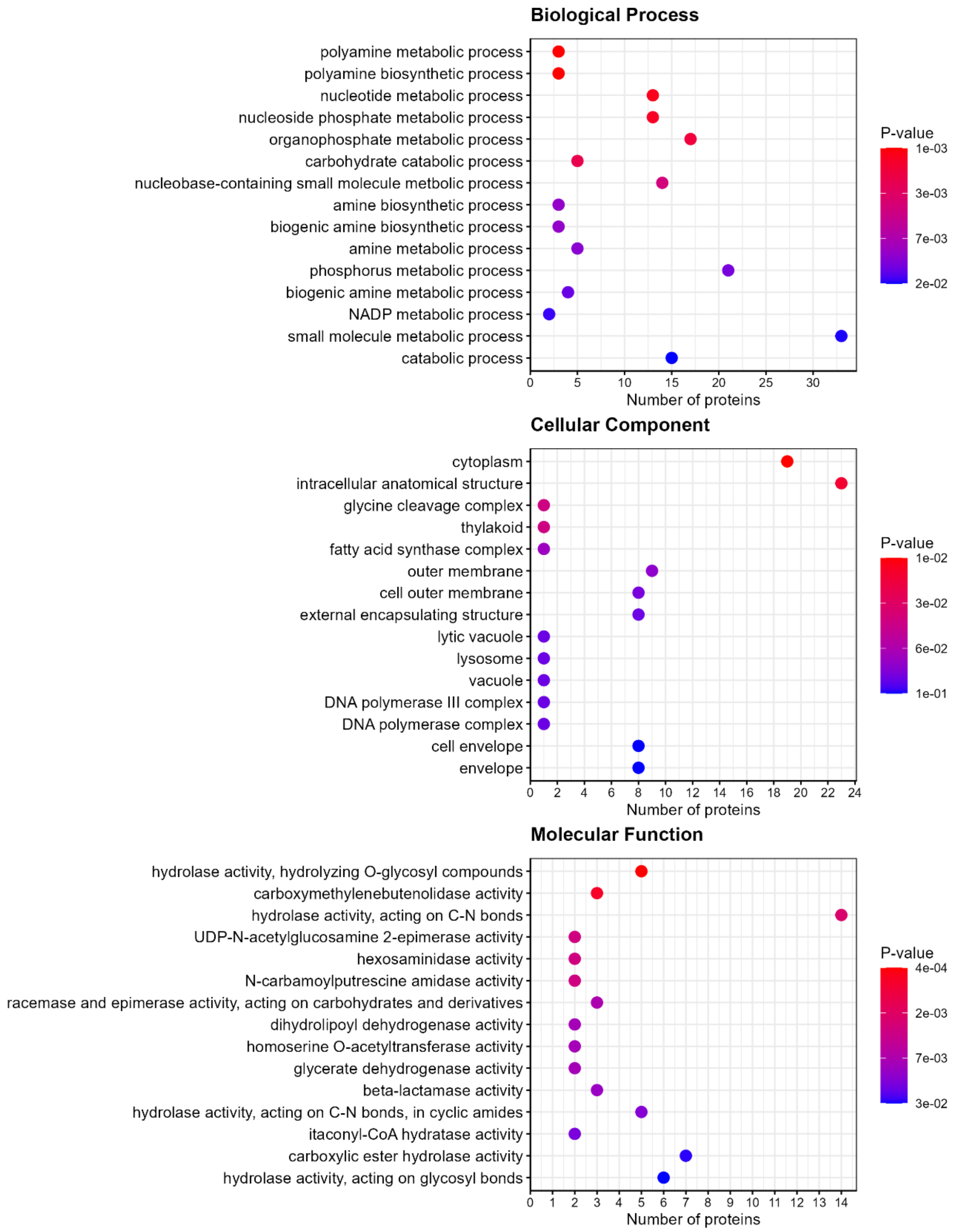


**Supplementary Figure 6: GO term enrichment of proteins downregulated in Δ*suhB* relative to the WT during stationary phase.** The X-axis shows the number of genes per category, and bar color indicates the p-value, with redder bars representing lower p-values.

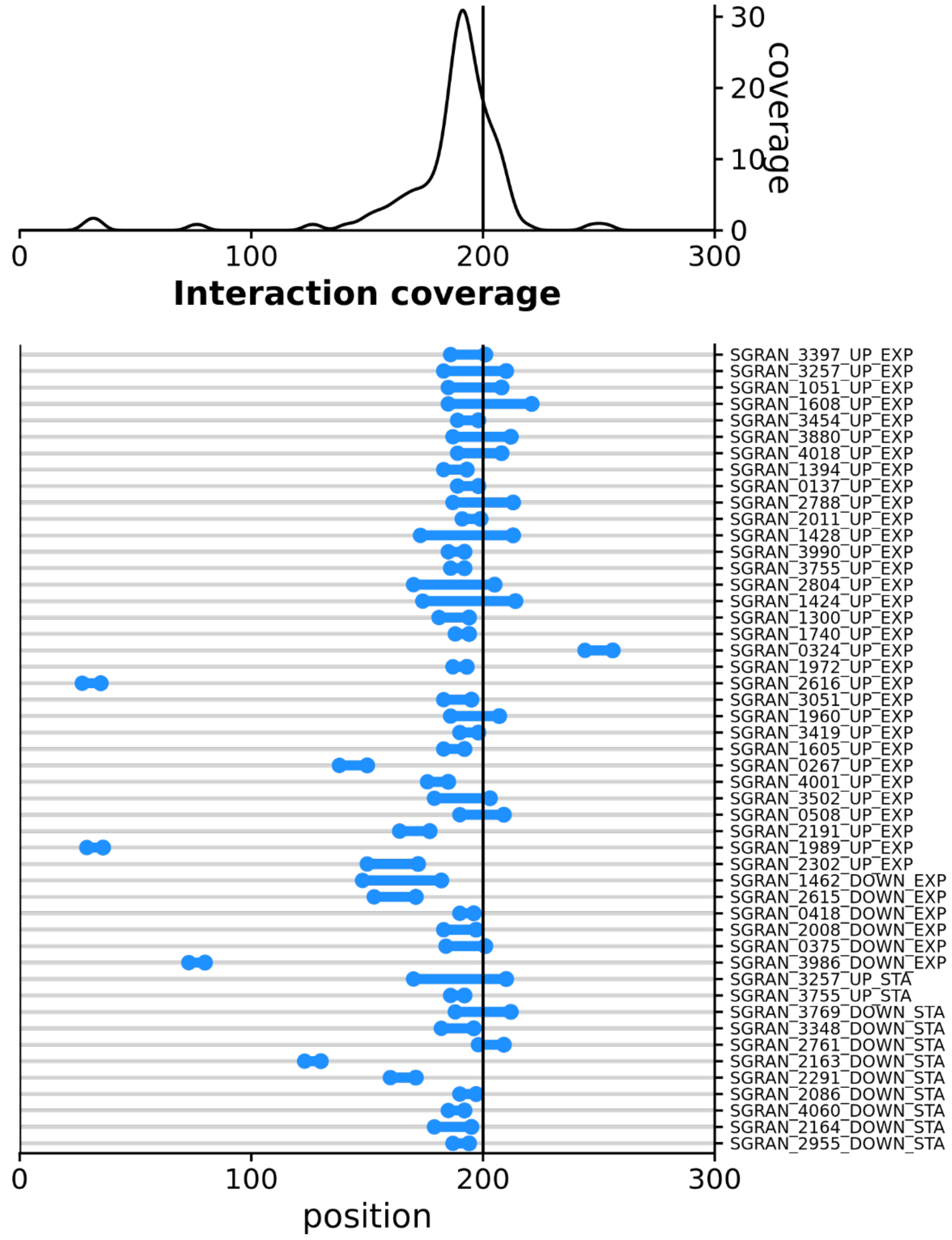


**Supplementary Figure 7: Coverage of the predicted mRNA-SuhB interactions across target sequences.** Each horizontal line represents a predicted interaction region, with the blue dots marking the start and end positions of the interactions. The vertical black line indicates the start codon. UP_EXT and DOWN_EXT denote proteins up or downregulated, respectively, in Δ*suhB* during exponential phase, while UP_STA and DOWN_STA denote up or downregulated proteins in stationary phase.

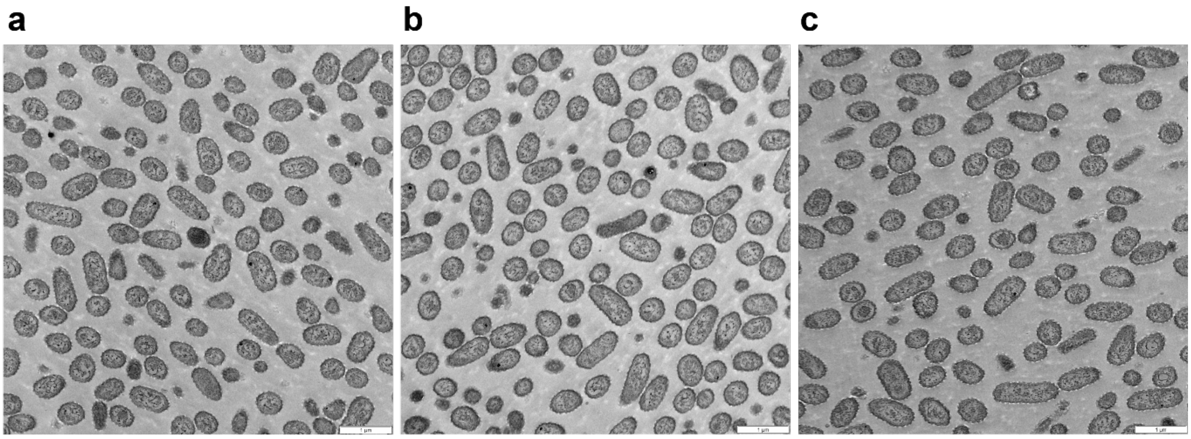

**Supplementary Figure 8: Transmission electron micrographs of cells in stationary phase.** (a) *Sphingopyxis granuli* TFA, WT; (b) Δ*suhB* mutant; (c) *miniTn5::phaC* mutant.

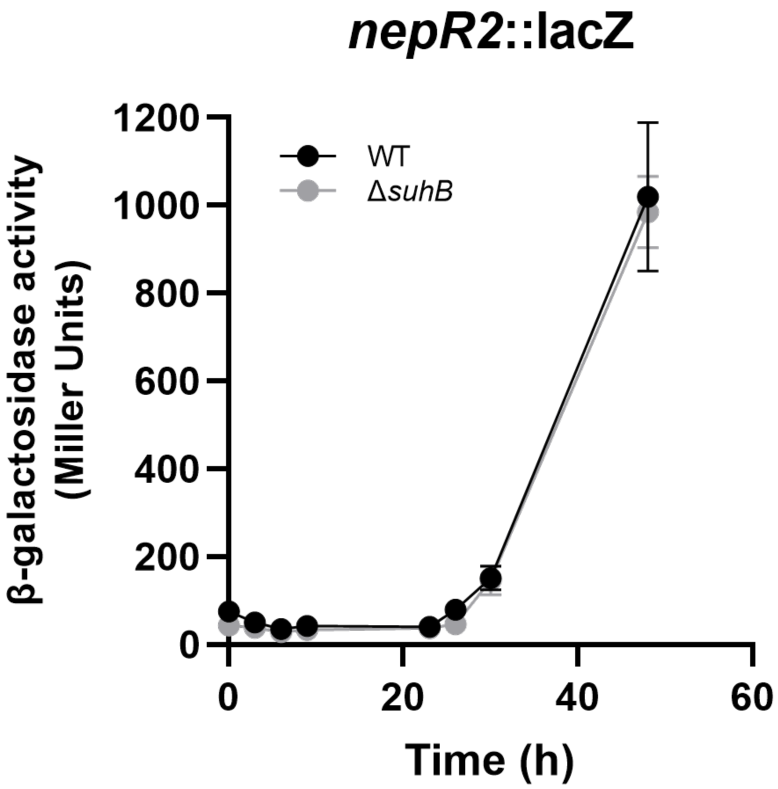


**Supplementary Figure 9: Expression of *nepR2* in WT and Δ*suhB*.** β-galactosidase activity from the *nepR2::lacZ* translational fusion in cells growing in minimal medium supplemented with 40 mM of BHB.

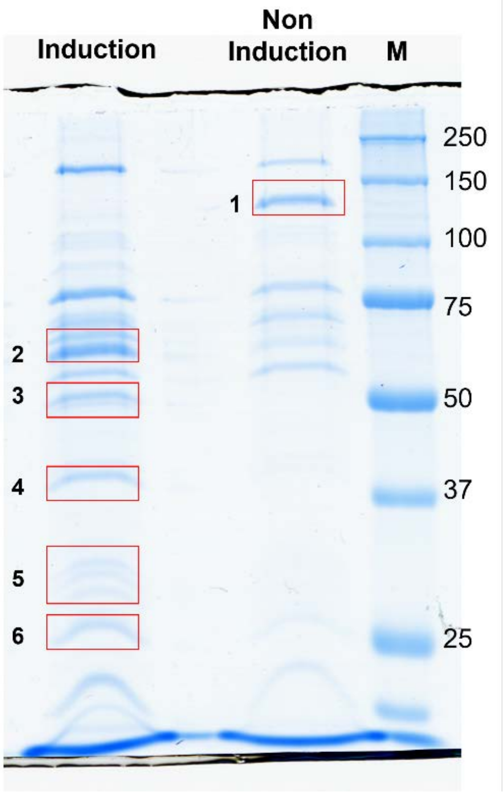


**Supplementary Figure 10: SDS-PAGE analysis of proteins bound to the *suhB* promoter.** Lane 1 shows the proteins bound to the *suhB* promoter isolated from cells grown in minimal medium supplemented with 16 mM sebacic acid. Lane 2 contains the proteins bound to the promoter obtained from cells grown in minimal medium supplemented with tetralin. Lane 3 represents the molecular weight marker, Precision Plus Protein™ Unstained Protein Standards (Bio-Rad). Numbers 1 to 6 correspond to protein bands that were subsequently analyzed by MALDI-TOF/TOF mass spectrometry.

### Supplementary References


1. D. Hanahan, Studies on transformation of Escherichia coli with plasmids. *Journal of Molecular Biology* **166**, 557-580 (1983).
2. X. Luo, N. Majdalani, Directed Screening for sRNA Targets in E. coli Using a Plasmid Library. *Methods Mol Biol* **2741**, 291-306 (2024).
3. M. J. Hernáez, W. Reineke, E. Santero, Genetic analysis of biodegradation of tetralin by a Sphingomonas strain. *Appl Environ Microbiol* **65**, 1806-1810 (1999).
4. G. Martín-Cabello, E. Moreno-Ruiz, V. Morales, B. Floriano, E. Santero, Involvement of poly(3-hydroxybutyrate) synthesis in catabolite repression of tetralin biodegradation genes in Sphingomonas macrogolitabida strain TFA. *Environ Microbiol Rep* **3**, 627-631 (2011).
5. I. García-Romero, K. U. Förstner, E. Santero, B. Floriano, SuhB, a small non-coding RNA involved in catabolite repression of tetralin degradation genes in Sphingopyxis granuli strain TFA. *Environ Microbiol* **20**, 3671-3683 (2018).
6. R. de Dios, E. Rivas-Marin, E. Santero, F. Reyes-Ramírez, Two paralogous EcfG σ factors hierarchically orchestrate the activation of the General Stress Response in Sphingopyxis granuli TFA. *Sci Rep* **10**, 5177 (2020).
7. B. Staskawicz, D. Dahlbeck, N. Keen, C. Napoli, Molecular characterization of cloned avirulence genes from race 0 and race 1 of Pseudomonas syringae pv. glycinea. *J Bacteriol* **169**, 5789-5794 (1987).
8. I. García-Romero, R. de Dios, F. Reyes-Ramírez, An improved genome editing system for Sphingomonadaceae. *Access Microbiol* **6**,  (2024).
9. R. Silva-Rocha *et al.*, The Standard European Vector Architecture (SEVA): a coherent platform for the analysis and deployment of complex prokaryotic phenotypes. *Nucleic Acids Res* **41**, D666-675 (2013).
10. E. Martínez-García, V. de Lorenzo, Engineering multiple genomic deletions in Gram-negative bacteria: analysis of the multi-resistant antibiotic profile of Pseudomonas putida KT2440. *Environ Microbiol* **13**, 2702-2716 (2011).
